# Striatum-projecting prefrontal cortex neurons support working memory maintenance

**DOI:** 10.1101/2021.12.03.471159

**Authors:** Maria Chernysheva, Yaroslav Sych, Aleksejs Fomins, José Luis Alatorre Warren, Christopher Lewis, Laia Serratosa Capdevila, Roman Boehringer, Elizabeth A. Amadei, Benjamin F. Grewe, Eoin C. O’Connor, Benjamin J. Hall, Fritjof Helmchen

## Abstract

The medial prefrontal cortex (mPFC) and the dorsomedial striatum (dmStr) are linked to working memory (WM) but how striatum-projecting mPFC neurons contribute to WM encoding, maintenance, or retrieval remains unclear. Here, we probed mPFC→dmStr pathway function in freely-moving mice during a T-maze alternation test of spatial WM. Fiber photometry of GCaMP6m-labeled mPFC→dmStr projection neurons revealed strongest activity during the delay period that requires WM maintenance. Demonstrating causality, optogenetic inhibition of mPFC→dmStr neurons only during the delay period impaired performance. Conversely, enhancing mPFC→dmStr pathway activity—via pharmacological suppression of HCN1 or by optogenetic activation during the delay— alleviated WM impairment induced by NMDA receptor blockade. Consistently, cellular-resolution miniscope imaging resolved preferred activation of >50% mPFC→dmStr neurons during WM maintenance. This subpopulation was distinct from neurons showing preference for encoding and retrieval. In all periods, including the delay, neuronal sequences were evident. Striatum-projecting mPFC neurons thus critically contribute to spatial WM maintenance.

## INTRODUCTION

Working memory (WM) is the ability to temporarily hold and manipulate relevant information in one’s memory to guide future actions, a process that is essential for many forms of goal-directed behaviour. The medial prefrontal cortex (mPFC), which is densely connected to many other brain regions, has a central function in executing tasks that require WM^1,2^. Indeed, dysfunctions of mPFC, as they occur in several mental disorders including schizophrenia, are associated with WM deficits^3–5^. Despite extensive research on mPFC, we still incompletely understand how mPFC neurons contribute to WM and what type of information they transfer via specific pathways to downstream target areas^6^.

Neurons in the mPFC receive inputs from a diverse set of brain structures and their major axonal projections target the mediodorsal nucleus (MD) in the thalamus, the dorsomedial striatum (dmStr), the basolateral amygdala (BLA), and the ventral tegmental area (VTA)^6^. Recent studies have begun to dissect the specific WM contributions of afferent and efferent pathways in this complex network. For example, projections from hippocampus to mPFC in mice were found to be critical for encoding, but not maintenance or retrieval, of spatial cues in a WM task^7^. In addition, analysis of mPFC interactions with MD in the thalamus revealed that activity of the MD→mPFC pathway is essential for sustaining prefrontal activity during WM maintenance^7–9^. The reciprocal mPFC→MD pathway, on the other hand, was found to mainly guide successful WM retrieval and to support subsequent choice^8^. This efferent pathway appears less important for WM maintenance during the delay period, when MD activity actually leads activity in the mPFC^8^, and it is not essential for encoding. A further dissection of the functional roles of specific pathways emerging from mPFC (or targeting mPFC) is needed to puzzle together the precise involvement of mPFC in WM, and especially WM maintenance.

Here, we hypothesize that the projection from mPFC to dmStr is a likely candidate pathway to support maintenance of information in a WM task. Consistent with this hypothesis, both the prelimbic (PrL) and infralimbic (IL) regions of mouse mPFC contain neurons that project to the anterior dmStr portion^10,11^. Moreover, lesions of dorsomedial, but not dorsolateral, striatum result in WM impairments^12,13^. Additionally, electrophysiological recordings from either mPFC or dmStr revealed neuronal populations that display activity patterns spanning WM maintenance periods in a sequential manner^8,14–16^. Therefore, we consider it likely that the mPFC→dmStr pathway is part of the brain circuitry that is essential for delayed goal-directed behaviour, albeit it remains unclear what exact information is conveyed between these two structures. To specifically probe the function of this pathway in a T-maze alternation task, we performed fiber-optic calcium recordings of mPFC→dmStr activity as well as cellular-resolution miniscope imaging to analyse activity patterns in striatum-projecting mPFC neurons. Furthermore, we tested the computational role of the mPFC→dmStr projection neurons during specific periods of the WM task using optogenetic and pharmacological manipulations. Our results corroborate the notion that the mPFC→dmStr pathway is critically involved in WM maintenance periods.

## RESULTS

### Fiber photometry of mPFC→dmStr pathway activity during T-maze alternation

We trained C57BL/6 mice in a T-maze alternation task commonly employed to study spatial WM^17–19^. In this task, freely moving mice receive a water reward during a ‘choice run’ when they correctly choose the left or right arm of the T-maze that is opposite to the arm they visited in the previous ‘sample run’ (Methods). Each trial consists of three periods related to the *encoding, maintenance*, and *retrieval* of WM (**Fig. 1a**). In the encoding period (sample run), one of the two T-maze arms is blocked by a door and the mouse is directed towards a first water reward in the open arm. After turning back towards the start box at the end of the sample run and throughout the subsequent delay period, the mouse must maintain information about the location of the sample reward in WM. In our setup, we enforced a delay waiting period in the start box of at least 5 seconds (5 s for fiber photometry and miniscope experiments; 10 s for optogenetic and pharmacological perturbations; Methods). Finally, in the choice run, all doors open and the mouse must retrieve the information held in WM in order to correctly alternate to the T-maze arm previously not visited. Correct choices result in a second water reward, triggered by licking at the water spout. Upon a mistake, no second reward is delivered and the mouse has to return to the start box to initialize the next trial. After completing training (Methods), mice typically reached performance levels of 70-80% correct trials (**Supplementary Fig. 1a**). Because WM is robustly impaired by the NMDA-type glutamate receptor (NMDAR)-blocker MK-801^20,21^, we tested in 11 mice whether such manipulation affected alternation performance under our conditions. Confirming previous results^17,22^, acute systemic MK-801 application (0.1 mg/kg) significantly reduced performance and shortened trial durations, across all periods, indicating hyperlocomotion (**Fig. 1b**). Importantly, mice still correctly completed the general task sequence, initiating licking on the water-spout and returning to the start box. The effect of MK-801 cannot be attributed to a specific neural pathway, though.

**Figure 1.**
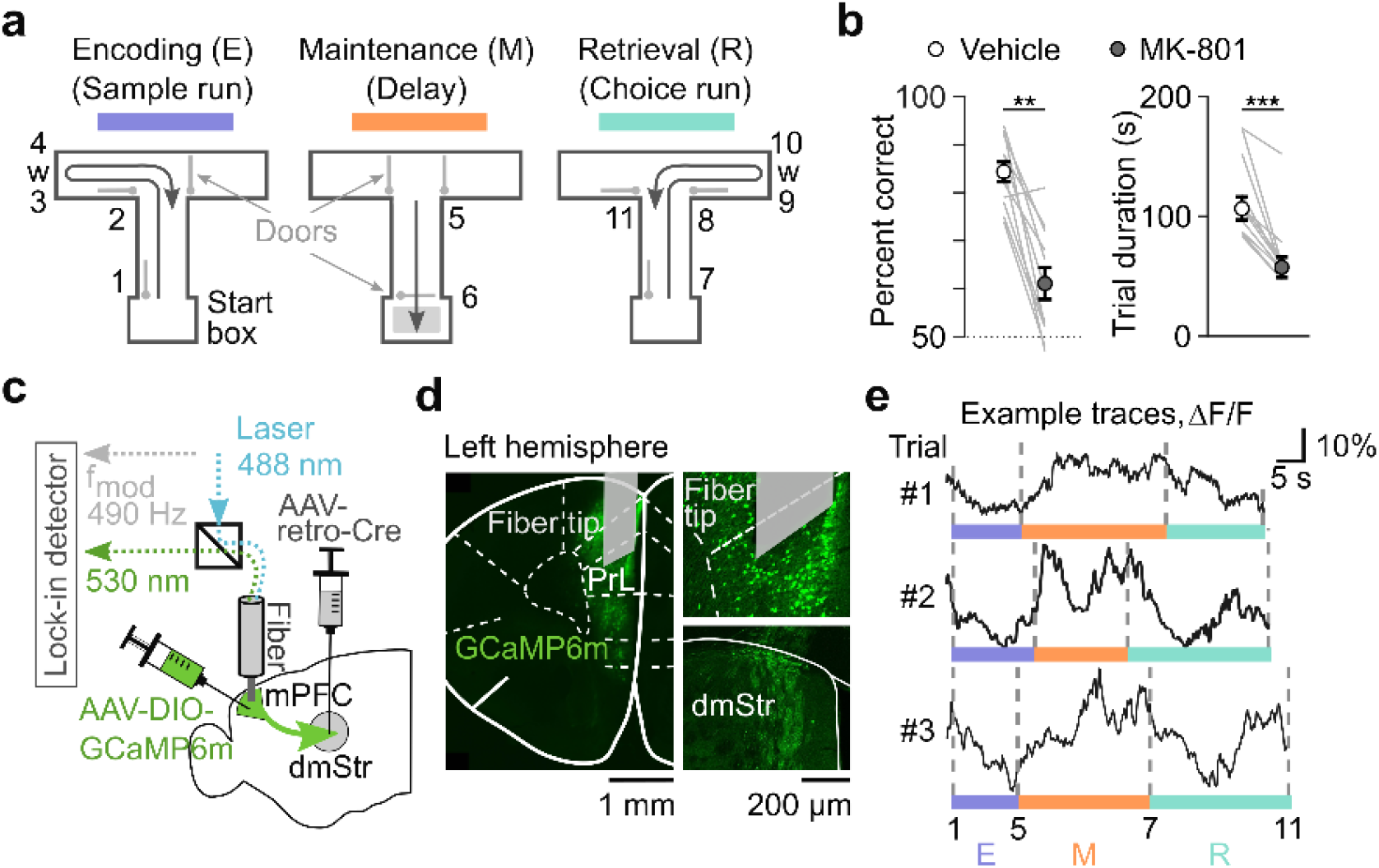
Fiber photometry of calcium signals in striatum-projecting mPFC neurons during T-maze alternation. (**a**) T-maze alternation task with automated doors to enforce sample runs and enable choice runs. Numbers indicate salient task events: 1–start of the sample run, 2–turning at the T-junction of the main maze arm, 3–first water reward, 4–end of licking period, 5–turn to run towards the start box, 6–reaching the start box, 7–start of the choice run, 8–turning at the T-junction, 9–second water reward, 10–end of licking period, 11–end of choice run. The second water reward is omitted in incorrect trials. Encoding (1 - 5), maintenance (5 - 7), and retrieval (7 - 11) periods are colored in purple, orange, and cyan, respectively. (**b**) Left: Task performance is impaired by the NMDAR-blocker MK-801 (0.1 mg/kg, i.p.). Right: MK-801 reduced mean trial duration significantly. n = 11 mice, mean ± s.e.m., ***p < 0.001, **p < 0.01, paired Wilcoxon signed rank test. (**c**) Schematic of virus injections and fiber implantation in the left hemisphere for photometric calcium recordings of dmStr-projecting mPFC neurons. (**d**) Left: Cre-dependent expression of GCaMP6m in a coronal histological section with implanted fiber tip shown above the prelimbic area (PrL). Right, top: Confocal images of GCaMP6m-labeled cell bodies in PrL (+1.8 AP, Methods) and bottom: axonal terminals in dmStr (+1.05 AP, Methods). (**e**) Example ΔF/F traces of pathway-specific photometric GCaMP6m recordings in PrL for 3 correct trials. Dashed vertical lines and purple, orange and cyan horizontal bars indicate the different task periods according to panel a. Note the variable duration of trial periods across trials.

To specifically measure the activity of mPFC→dmStr projection neurons during alternation behaviour, we injected retrograde Cre-expressing virus unilaterally into the left dmStr and a Cre-dependent GCaMP6m-expressing virus into left mPFC, targeting the prelimbic area PrL (n = 6 mice; Methods). To perform fiber photometry, we chronically implanted an optical fiber above the mPFC injection site (**Fig. 1c,d**). Once mice had reached expert level (≥60% correct trials on 2 consecutive days), we performed pathway-specific calcium recordings while mice executed the alternation task. Relative percentage changes in fluorescence (ΔF/F) showed trial-related activity with transient changes up to 30% amplitude and variable time courses across trials and mice, indicating spiking activity of mPFC→dmStr projection neurons (**Fig. 1e**). These ΔF/F traces were not confounded by motion artefacts as demonstrated by control ΔF/F signals excited at 425-nm wavelength (at which GCaMP6m has low calcium-sensitivity) that displayed substantially lower variation compared to 488-nm excited signals (**Supplementary Fig. 1b,c**). Taken together, these experimental settings enabled us to study WM-related activity in the mPFC→dmStr pathway.

### The mPFC→dmStr pathway sustains high activity during WM maintenance

Based on 11 salient trial-related events we defined 10 trial phases (**Fig. 1a**). Because mice behaved freely, the duration of these phases varied from trial to trial, including the WM maintenance period between turning back towards the start box (event 5) and the start of the choice run (event 7) (**Fig. 1e**). The mean durations of encoding, maintenance and retrieval periods did not differ, however, for correct choices and mistakes (**Supplementary Fig. 1d**). To analyse the temporal profile of mPFC→dmStr activity across trials, sessions, and animals we had to account for the variability of trial phase duration. To this end, we aligned the photometric calcium signals to the 10 salient trial phases by segment-wise resampling the ΔF/F traces so that each phase duration matched the median of the corresponding phase duration across all trials (Methods). We then averaged the resampled and z-scored ΔF/F traces for 6 GCaMPm-expressing mice and 4 GFP-expressing control mice (n = 5 sessions for each mouse). The mPFC→dmStr pathway exhibited the highest activity during the WM period, with a strong signal increase at the beginning of the return to the start box. The signal remained high, albeit slowly declining, until the start of the choice run (**Fig. 2a**). The mean activity in the maintenance period was significantly higher compared to both retrieval and encoding period (**Fig. 2b**; p < 0.001, Fisher’s method). In control GFP mice, the average ΔF/F trace was flat with no difference in mean activity between trial periods (**Fig. 2b**).

**Figure 2.**
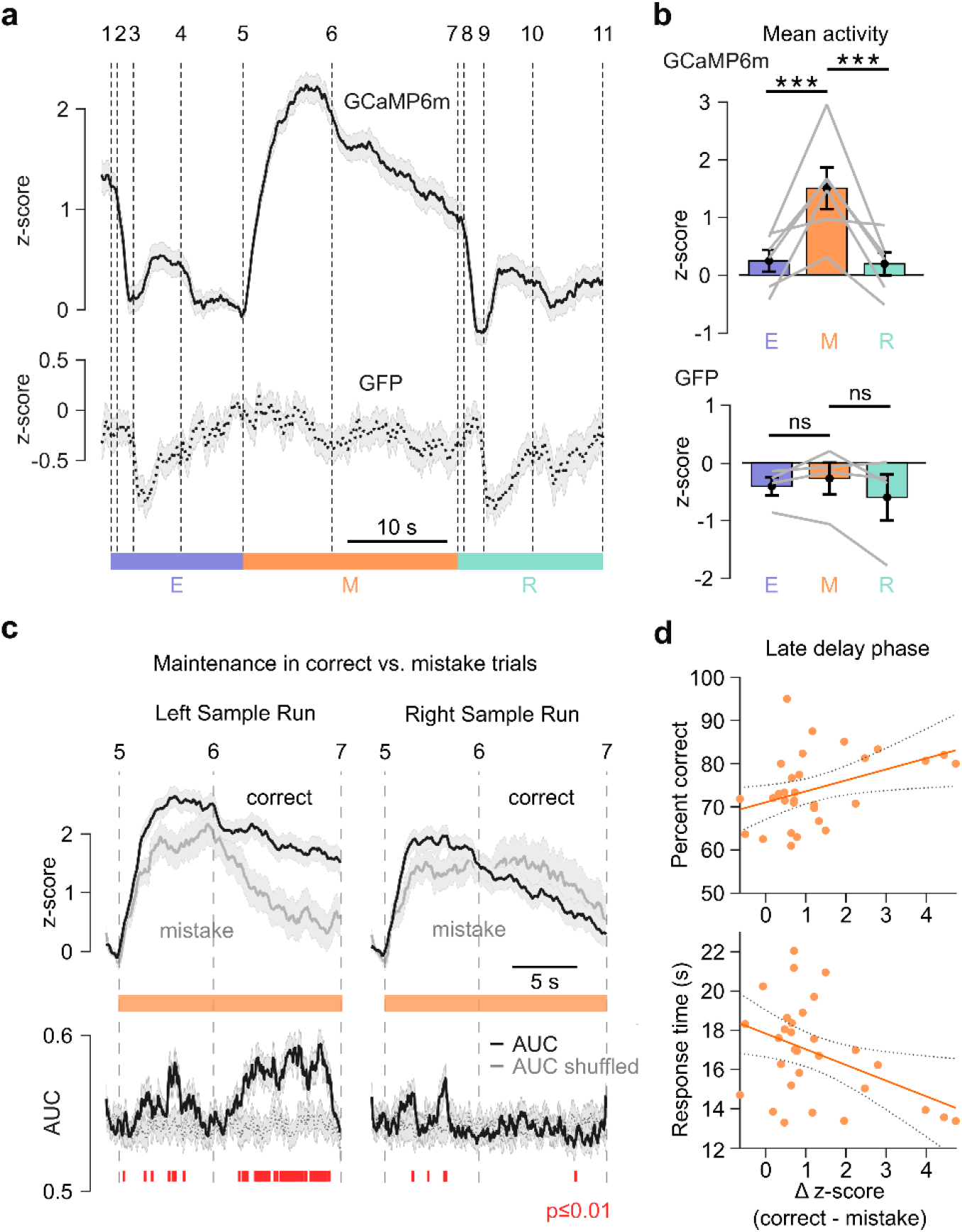
mPFC→mStr activity correlates with WM maintenance. (**a**) Resampled and session-averaged z-scored calcium signal (n = 6 mice; solid line) and non-calcium dependent GFP control signal (n = 4 mice; dashed line). Vertical dashed lines and numbers indicate trial events. Shaded areas indicate s.e.m. (**b**) Mean z-score values over the different task epochs. Each grey line represents one mouse (***p < 0.001, p-values combined for mice using Fisher’s method; n = 6 mice; 103-156 correct trials per mouse; paired Wilcoxon signed test performed for individual mice). (**c**) Top: Mean z-scored calcium signals during the maintenance period for correct alternation in choice runs (black) and for mistakes (grey). Data are shown separately for periods following left sample runs (correct choice to the right, contra-lateral to the recording site) and right sample runs (correct choice to the ipsi-lateral side). Bottom: Time course of correct vs. mistake classification based on the area-under-the-curve (AUC) of an ROC analysis. Red markers at the bottom indicate time bins with significant classification accuracy compared to trial-shuffled data (grey trace; p < 0.01; Mann-Whitney U-test). (**d**) Top: Task performance positively correlates with the difference in mean z-scores (Δz-score, left sample correct trials minus left mistake) for the late delay period (n = 32 sessions, R-squared = 0.163, p = 0.026, linear regression model). Bottom: Response time (defined as time from opening of the start box to detecting the mouse in the chosen arm) negatively correlated with Δz-score (n = 32 sessions, R-squared=0.176, p=0.021), indicating that the mouse made its decision with higher confidence for increased Δz-scores. Error bars show s.e.m.

We further assessed whether mPFC→dmStr activity differs for correct versus mistake trials. Because the activity of mPFC neurons in one brain hemisphere may depend on the location of the choice reward (left versus right goal arm)^16,23^, we further stratified the fluorescence signals—recorded in the left hemisphere—into contra-lateral (right) and ipsilateral (left) choices. Indeed, fluorescence signals during the maintenance period were larger for correct choices towards the contralateral side compared to correct ipsilateral choices (**Supplementary Fig. 2**). For contralateral choice runs, mPFC→dmStr activity was significantly enhanced in the maintenance period when the animal’s turn was correct compared to mistake trials (**Fig. 2c**). These results are in line with previous findings that the activation of striatal neurons in one hemisphere precedes the initiation of contralateral movements^24^. The most significant difference between correct and mistake trials occurred in the late maintenance phase for contralateral choices (**Fig. 2c**; second half of the phase between events 6 and 7; quantified by a receiver operating characteristics (ROC) analysis; Methods). In this time window, the level of mPFC→dmStr activity positively correlated with task performance and negatively correlated with response time in correct trials, which indicates the confidence with which mice make their decision (**Fig. 2d**). These findings suggest that the performance in the alternation task may critically depend on the mPFC→dmStr pathway activity specifically during the WM maintenance period.

### mPFC→dmStr pathway activity is required for WM maintenance

Guided by our photometry results, we next aimed to test whether pathway-specific optogenetic perturbation of neural activity during the maintenance period would lead to changes in task-related behavior. First, we tested if mPFC→dmStr activity is required for WM maintenance using optogenetic silencing. To selectively inhibit this pathway, we injected retrograde AAV-Cre into the left dmStr and Cre-dependent AAV driving expression of the light-driven proton pump archaerhodopsin ArchT^25^ into the left PrL. This approach resulted in strong ArchT expression in striatum-projecting PFC neurons (**Fig. 3a**; n = 7 mice; Methods). ArchT has been previously applied to silence mPFC neurons^7,8,26,27^ but for further validation we verified in 7 ArchT-expressing mice that 560-nm illumination indeed reduced multi-unit activity in both mPFC and downstream dmStr (**Fig. 3b,c**; **Supplementary Fig. 3**). In expert mice, we then transiently suppressed mPFC→dmStr activity during task performance by temporally restricting laser illumination to one of the three WM periods (encoding, maintenance, or retrieval). In each session, green light was delivered in 50% of randomly interleaved trials and targeted to only one of the WM periods. Optogenetic silencing during the maintenance period resulted in a significant performance decrease of 9% (**Fig. 3d**; p = 0.0035, paired Wilcoxon signed-rank test), whereas no significant behavioural effect was induced by silencing during the encoding or retrieval period. Illumination during the maintenance period did not affect the mean duration of choice runs (**Fig. 3e**), indicating that the performance decrease was not simply induced by altered locomotion. These findings are consistent with the results of our photometry recordings and corroborate the notion that mPFC→dmStr pathway activity is required for WM maintenance.

**Figure 3.**
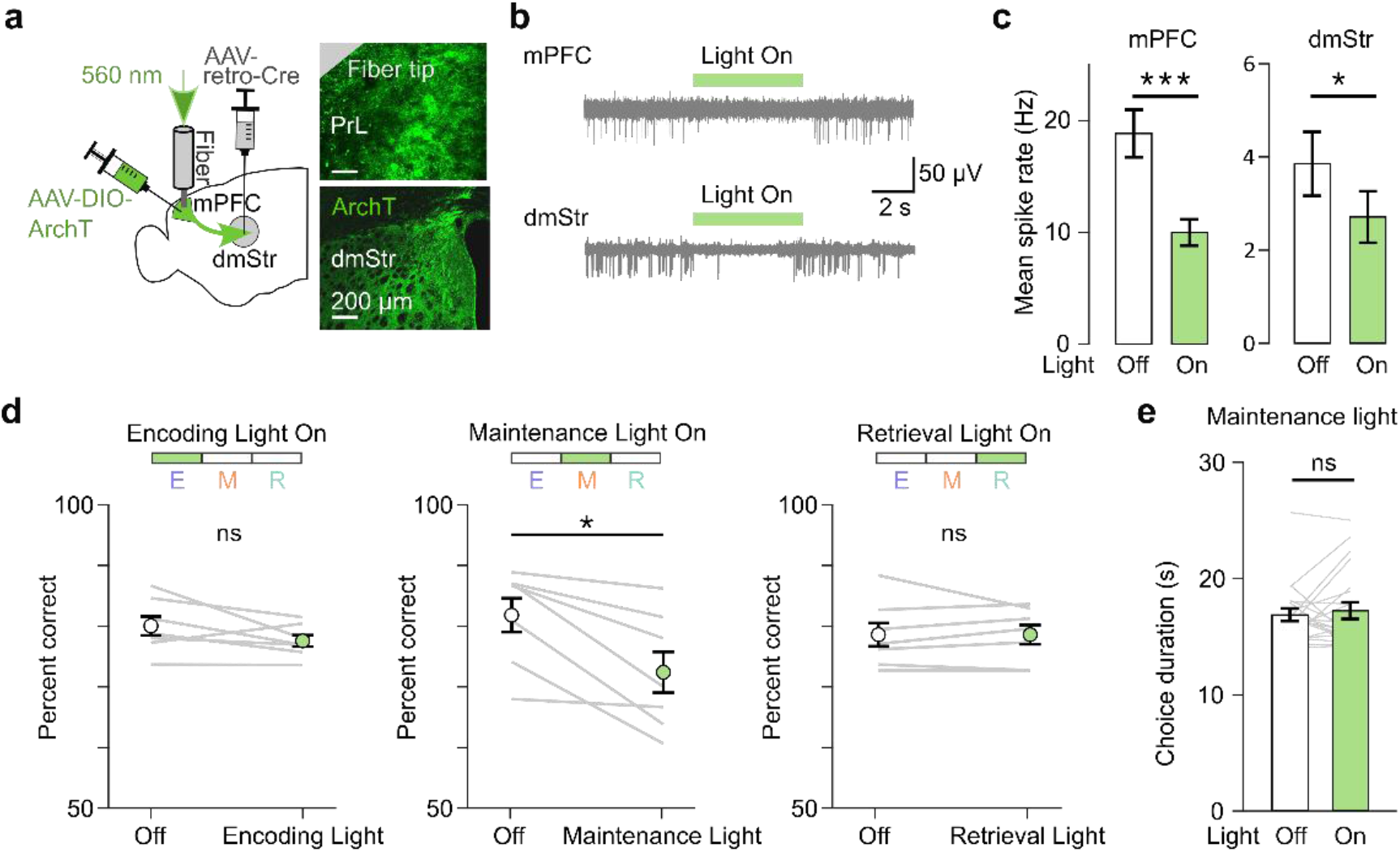
Photoinhibition of mPFC→dmStr activity during WM maintenance impairs task performance. (**a**) Left: Schematic diagram of viral injections and optical fiber implantation in the left hemisphere allowing for pathway-specific ArchT expression and photoinhibition of mPFC→dmStr projection neurons. Right: confocal images of ArchT-expressing cells in PrL below the fiber tip (top) and ArchT-expressing axons in dmStr (bottom). Example of multi-unit recordings from cells in mPFC (top) and dmStr (bottom) showing suppressed activity during 560-nm illumination of ArchT-expressing mPFC neurons (indicated by green bars). (**c**) Light-induced suppression of multi-unit activity in mPFC and dmStr (***p < 0.001, 40 units from n = 2 mice; *p < 0.05, 10 units from n = 2 mice; Mann-Whitney U-test). (**d**) Performance changes upon silencing mPFC→dmStr projection neurons during different task periods. For each period, photoinhibition was enabled in random 50% of trials. Grey lines represent individual mice (n = 7 mice). Mean ± s.e.m. *p < 0.05, paired Wilcoxon signed-rank test. All other comparisons non-significant (p > 0.1). (**e**) Lack of effect of photoinhibition on the choice duration. Each line represents average choice duration (both contra- and ipsilateral turn trials) during individual sessions (p = 0.88, n = 7 mice, 3 sessions per mouse).

### mPFC→dmStr pathway activation alleviates WM impairment induced by MK-801

Given that elevated mPFC→dmStr activity positively correlates with task performance and that silencing of this pathway impairs WM, we next sought to establish whether enhancing mPFC→dmStr activity can improve performance or alleviate WM deficits. In a first approach, we assessed whether striatum-projecting neurons are affected by pharmacological blockade of HCN1 channels, an intervention that has been found to improve WM by enhancing mPFC neuronal activity during WM delay periods^28–30^. Here, we applied the HCN-channel blocker J&J12e that readily passes the blood-brain barrier^31^. After training mice in the alternation task with a 5-s delay, we tested performance with a challenging 10-s delay after systemic injection of either J&J12e or vehicle. Photometry revealed a significant enhancement of mPFC→dmStr activity after mice received the J&J12e (**Fig. 4a; Supplementary Fig. 4**), indicating that HCN blockade may indeed increase WM-related activity of mPFC→dmStr projection neurons. Despite the elevated mPFC→dmStr activity that we observed during the maintenance period in J&J12e-treated mice, these mice did not show improved performance (**Fig. 4b**). Hence, to test whether high baseline performance may have masked a potential compound effect, we also applied J&J12e after impairing WM by systemic injection of MK-801. Indeed, co-administration of J&J12e alleviated the WM impairment induced by MK-801 (**Fig. 4c**; **Supplementary Fig. 4**).

**Figure 4.**
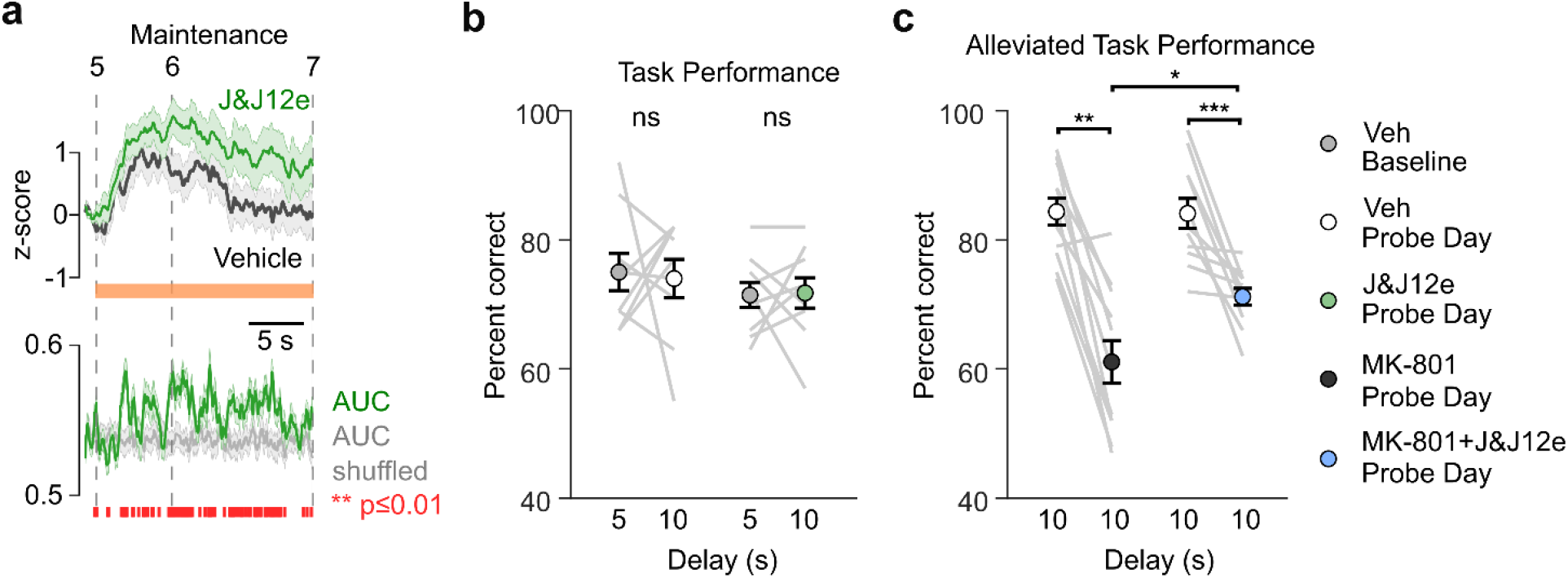
Blockade of HCN channels enhances WM-related mPFC→dmStr activity and rescues WM impairment induced by MK-801. (**a**) Pathway-specific activity is higher when mice receive the HCN-channel blocker J&J12e (green, 3 mg/kg, i.p.) compared to vehicle injection (grey). Average of z-scored ΔF/F traces for all correct trials (n = 6 mice). Bottom: Time course of classification power showing AUC of an ROC analysis. Red markers below indicate time bins with significant classification accuracy compared to trial-shuffled data (grey line; p < 0.01; Mann-Whitney U-test). Shaded areas indicate s.e.m. (**b**) No effect on performance was observed for both 5- s and 10-s delay conditions. (**c**) Impairment of alternation performance by NMDAR blockade with MK-801 (left, 0.1 mg/kg, i.p.) was alleviated by additional administration of J&J12e (right) as indicated by a smaller drop in percentage of correct trials. Grey lines represent individual mice (n = 11); error bars are s.e.m. ***p < 0.001, **p < 0.01, *p < 0.05, paired Wilcoxon signed rank test.

Systemic injection of an HCN-blocker presumably has broad and nonspecific effects on various brain regions. To test more specifically whether enhancing mPFC→dmStr pathway activity can improve task performance, we additionally took a second approach by inducing pathway-specific expression of channelrhodopsin-2 (ChR2) (**Fig. 5a**). We validated that transient continuous blue light illumination for ChR2 activation indeed induced action potentials in PrL neurons (**Fig. 5b** and **Supplementary Fig. 5**). ChR2 activation during the encoding and retrieval periods had no significant behavioural effect, whereas illumination during the maintenance period resulted in only a small performance increase of about 3% (**Fig. 5c;** illumination in GFP control mice had no significant effect; **Supplementary Fig. 6**). Again we reasoned that the high (∼80%) performance of untreated expert mice may have masked any further improvement. We therefore repeated ChR2-mediated mPFC→dmStr pathway activation after impairing WM by MK-801 application, lowering the initial performance levels to below 70%. Under these conditions, ChR2 activation significantly increased performance, but only if applied during the WM maintenance period (**Fig. 5d**). Optogenetic activation targeted specifically to the mPFC→dmStr pathway therefore had a similar effect of alleviating MK-801-induced WM impairment as the systemic administration of an HCN blocker.

**Figure 5.**
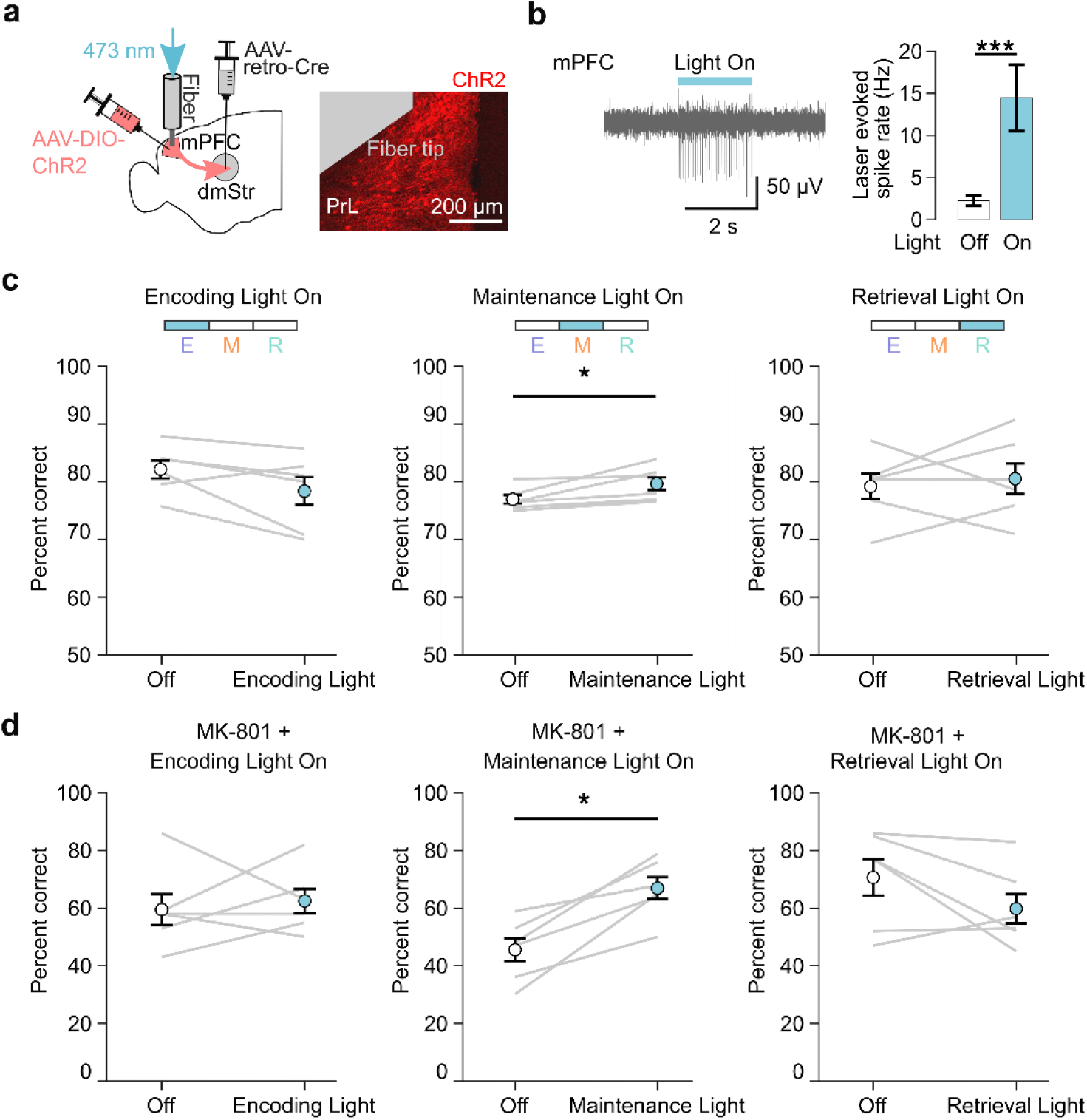
WM effects of optogenetic activation of the mPFC→dmStr pathway. (**a**) Virus injections to induce ChR2 expression in the mPFC→dmStr pathway and fiber implantation site in the left hemisphere. (**b**) Validation of pathway-specific activation induced by ChR2 activation. Left: example electrophysiological trace from a single unit in PrL; Right: mean evoked spike rate (32 units, from n = 2 mice) without (grey bar) and with (blue bar) 473-nm light illumination. (**c**) Optogenetic activation of the mPFC→dmStr pathway targeted to the three WM periods in Cre-dependent ChR2-expressing mice (n = 6 mice) induced a small performance change when applied in the WM maintenance period (77.0 ± 0.7% to 79.7 ± 1.1%; p = 0.03). (**d**) Optogenetic activation of the mPFC→dmStr pathway partially rescued WM impairment induced by MK-801 injection when applied in the WM maintenance period (performance increase from 45.5 ± 4.0% to 67.0 ± 3.8%; p = 0.03). For a given task period, ChR2 activation was performed in 50% of trials in a random fashion. Grey lines represent individual sessions. *p < 0.05, paired Wilcoxon signed-rank test. Error bars show s.e.m.

### Sequential activity in subpopulations of mPFC→dmStr projection neurons

Given that the mPFC→dmStr pathway is most active during the WM maintenance period and because optogenetic manipulations during this period affect behavior, we next asked whether a specific subpopulation of mPFC neurons might exhibit WM-related activity. To achieve cellular resolution we used a wearable miniaturized microscope (miniscope)^32,33^ and imaged mPFC neurons through a GRIN lens inserted into a chronically implanted cannula (**Fig. 6a**; Methods). We specifically labelled mPFC→dmStr projection neurons with GCaMP6f, employing the same dual-virus strategy as for fiber photometry in two mice. In addition, we achieved pathway-specific GCaMP6f expression by injecting retrograde Cre-expressing virus into dmStr of four transgenic Ai148D mice (**Fig. 6a**). Because these two labelling approaches yielded similar results, we pooled data across all 6 mice for analysis. Typically, we identified 30-90 longitudinally active neurons within a field of view (**Fig. 6b**; Methods).

**Figure 6.**
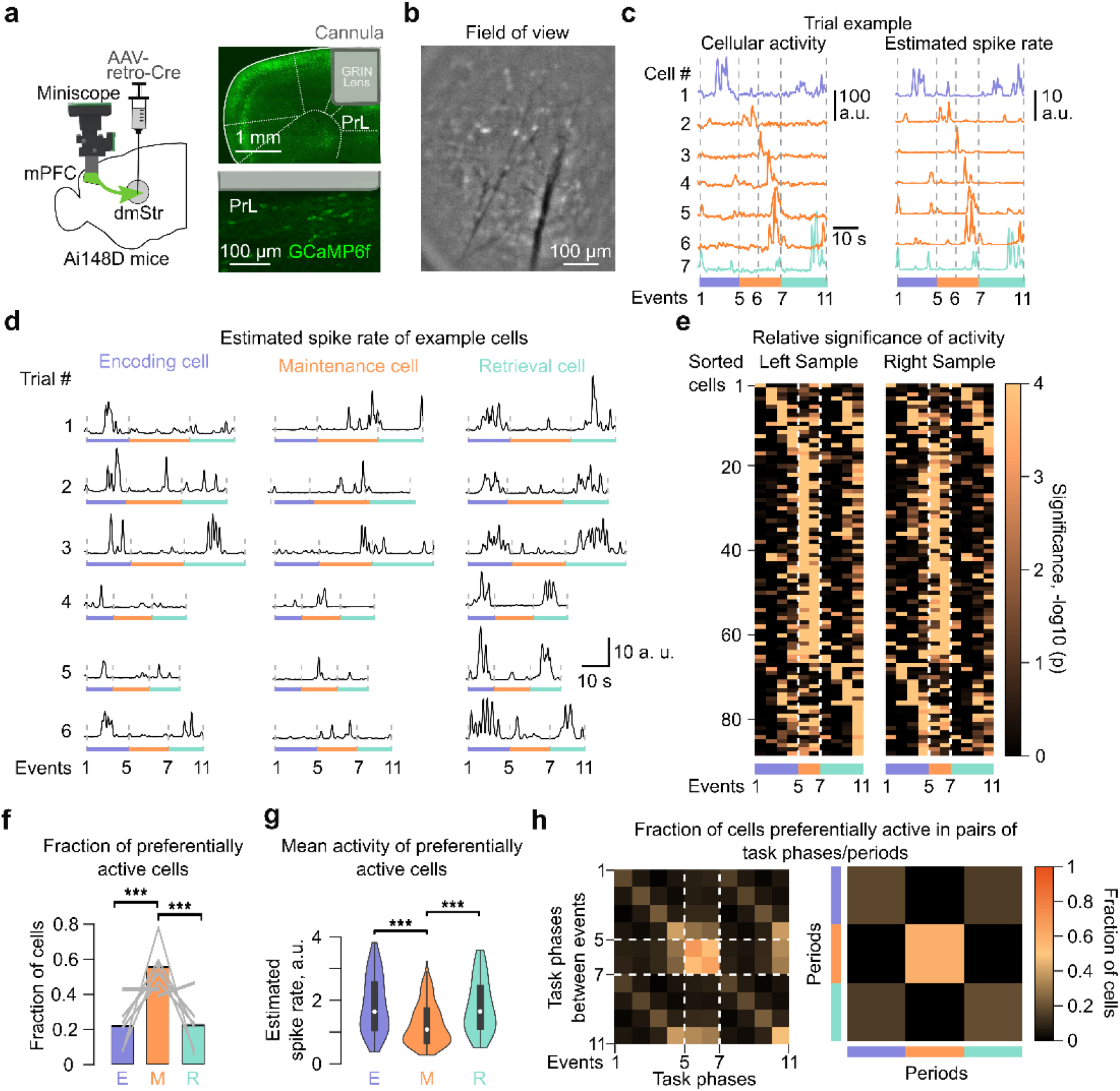
Miniscope calcium imaging of mPFC→dmStr projection neurons during T-maze alternation task. (**a**) Left: Schematic of the virus injection and miniscope mounting for cellular resolution imaging of mPFC→dmStr projection neurons. Right: Confocal images of GCaMP6f expression in left PrL indicating GRIN lens insertion into an implanted cannula. (**b**) Representative field of view for miniscope imaging (single time frame of pre-processed data). (**c**) Trial-related cellular activity traces of 7 example neurons for one correct trial. Left: output signal in arbitrary units, right: corresponding deconvolved estimated spike rate. Dashed lines, purple, orange and cyan bars indicate different task periods. (**d**) Example cells identified as being significantly more active in either encoding (left), maintenance (middle), or retrieval (right) period. Six example correct trials are plotted for each neuron. (**e**) Relative significance of single-cell activity during the 10 trial phases. High values indicate significantly elevated activity of a given cell in a particular trial phase, compared to the rest of the phases. The heatmap shows all cells for one example mouse. Only correct trials were analysed and cells sorted according to the Left Sample trials. (**f**) Fraction of cells preferentially active in encoding, maintenance, and retrieval period (***p<0.001, binomial test, p-values composed over mice using Fisher’s method; n = 6 mice). (**g**) Mean activity levels of neurons preferentially active in encoding (E), maintenance (M), and retrieval (R) period. Mean spike rate for a given period was calculated in a.u. for each trial and averaged for each neuron across all correct trials (n = 6 mice). (**h**) Matrix plot showing the fraction of neurons significantly active in pairs of task phases (left) or periods (right) (n = 6 mice). Values on the diagonal represent fraction of neurons significantly active in each respective phase or period. Note the overlap of neuronal fractions that show significant activity in encoding and retrieval periods as well as in corresponding phases during these periods.

We observed large task-related calcium transients, with example neurons showing preferential activation in distinct task periods (**Fig. 6c**). To test for consistency with the photometry experiments, we averaged task-related activity across all active mPFC→dmStr neurons within each FOV. These ‘bulk’ activity signals resembled the photometric signals, again revealing highest signal amplitude during the maintenance period as well as significantly elevated activity during the late delay period in correct versus mistake contralateral choice runs (**Supplementary Fig. 7**). For a more accurate representation of the time course of single-cell activities, we deconvolved neuronal ΔF/F traces using a novel spike inference algorithm^34^ (**Fig. 6c**). All subsequent analysis used the deconvolved traces (expressed in arbitrary units, a.u., because calibration in terms of absolute spike rates is uncertain). First, we analysed how individual neuron activity relates to the task phases as defined in Figure 1. Whereas some example neurons showed preferential activation during specific phases of the encoding or retrieval period, others were most active during the maintenance period (**Fig. 6d**).

To quantify the notion of preferential activity in distinct task periods, we calculated for each active neuron the significance level of it being active in each of the 10 trial phases (90-238 trials per mouse; Methods). Sorting neurons according to their most significant phase revealed that distinct subpopulations of neurons were preferentially active during the maintenance period and during the encoding/retrieval periods, respectively (**Fig. 6e** for an example mouse; **Supplementary Fig. 8** for all mice). The highest fraction of active mPFC→dmStr neurons occurred in the maintenance period (**Fig. 6f**) but the average activity per neuron was slightly lower during maintenance compared to encoding and retrieval (**Fig. 6g**). Thus, the large ‘bulk’ signals during maintenance predominantly reflect the relatively strong recruitment of mPFC→dmStr neurons during this task period.

We further quantified the differences between neuronal subpopulations by calculating the fraction of cells that were co-active in pairs of distinct task phases or periods (**Fig. 6h** for an example mouse; **Supplementary Fig. 9** for all mice). Populations preferentially active in encoding and retrieval periods showed substantial overlap, whereas the subpopulation preferentially active during the maintenance period was clearly distinct. In some mice, however, the populations preferentially active during encoding and retrieval also clearly reflected preference for either right or left turns (**Supplementary Fig. 9**). In addition, within the encoding and retrieval periods, different sets of neurons preferred distinct task phases, indicating sequences of neuronal activation during both sample and choice runs.

Previous studies also found neural sequences specifically during the delay period of WM tasks in both mPFC^8^ and dmStr^14^. Such sequences could help to maintain information in short-term memory. Therefore, we further evaluated the temporal order of the activity of mPFC→dmStr projection neurons during the maintenance period (phases 5-6 and 6-7). Some cells were consistently active at the beginning or at the end of the maintenance period whereas others displayed more variable and temporally distributed activity across trials (**Supplementary Fig. 10**). Overall, cells spanned the entire maintenance period with their activity (**Fig. 7a,b** for an example mouse; **Supplementary Fig. 11** for all cells across mice). We tested for consistency and significance by splitting all correct trials into half, creating a ‘training’ and a ‘test’ set. We then ordered neurons according to the peak times of the mean signals across the training set and applied the same ordering to the ‘test’ set as well as to the mean ΔF/F signals of mistake trials. Whereas the temporal order of neuronal activation was similar in test and training data for correct trials, the sequence of activity deteriorated for mistake trials (**Fig. 7b**; **Supplementary Fig. 11**) with response peaks shifting significantly further apart from training data compared to test trials (**Fig. 7c**).

**Figure 7.**
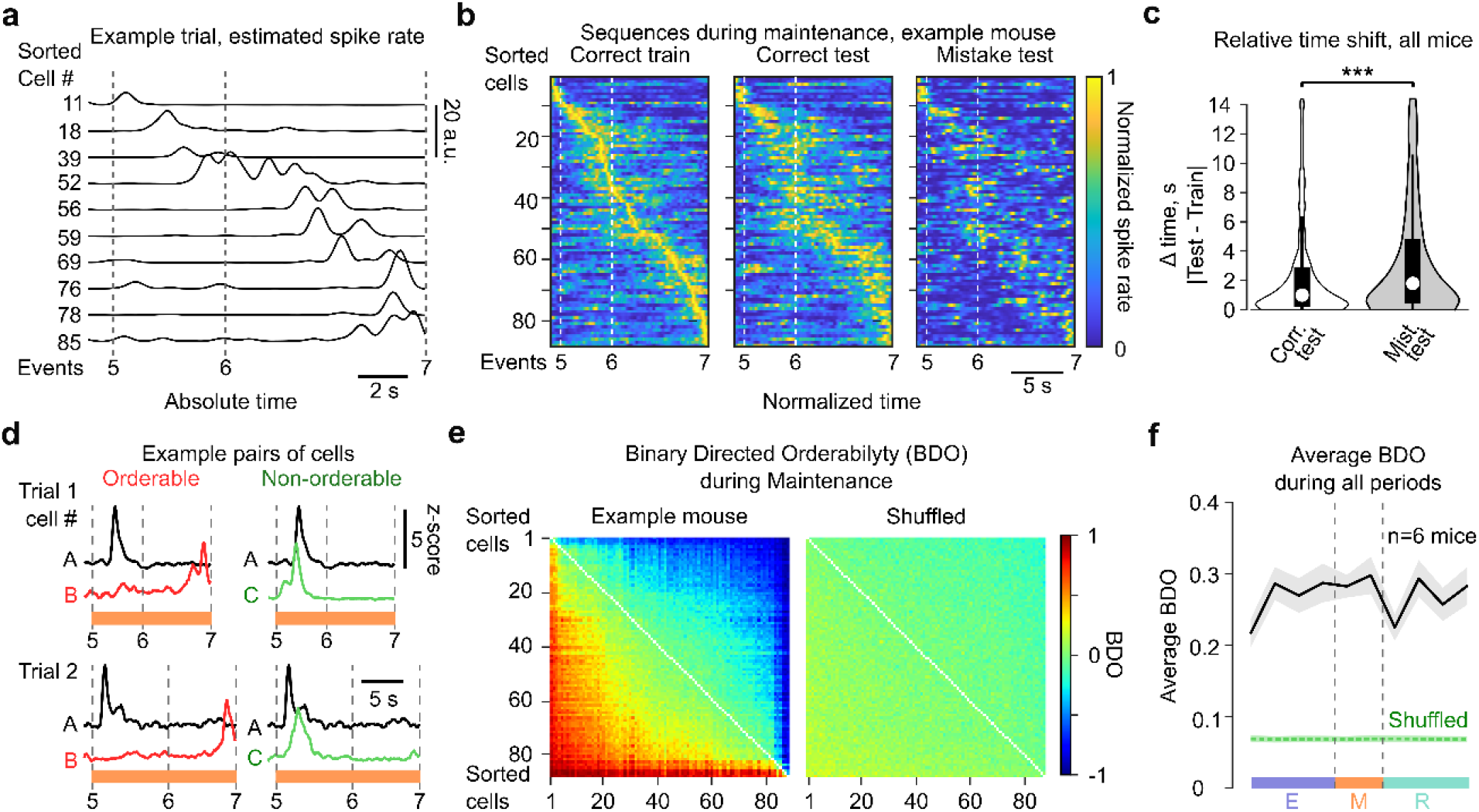
Sequential activation of PFC→dmStr neurons. **(a)** Representative deconvolved ΔF/F traces from 10 example maintenance cells. Cell numbers correspond to the sorting in b. **(b)** Normalized deconvolved neuronal signals during the maintenance period for an example mouse. Left: Mean signals for a randomly selected half of correct trials used as a training set. Middle: Mean signals for the remaining test set of correct trials. Right: Mean signals for mistake trials. In all plots, neurons are sorted according to the signal peak times in the training data. (**c**) Relative time shift of response peaks compared to training data for all neurons. Peak times shifted less for test trials (median = 1.0 s, 75th percentile = 2.8 s) compared to mistake trials (median = 1.80 s, 75th percentile = 4.8 s). ***p<0.001, paired Wilcoxon signed-rank test. (**d**) Examples of two trials, showing an orderable cell pair [A,B] (left) and a non-orderable pair [A,C] (right). Cell A is active earlier than cell B in most of the trials (BDO = 0.96), whereas cell C is active earlier than cell A in some trials but later in others, resulting in a lower BDO of 0.66. (**e**) Left: BDO matrix calculated for the maintenance period for all neurons recorded in an example mouse. Right: BDO matrix for shuffled data. Neurons are sorted by neuron-average BDO.(**f**) Average absolute value BDO (± s.e.m.) across all neuronal pairs for 6 mice (black), calculated for all 10 task phases and compared to shuffled data (green).

To further corroborate the notion of sequential activity, we also accounted for the observed trial-by-trial variability and evaluated whether neuronal pairs showed temporally ordered activation consistently across trials. To this end, we defined a novel ‘binary directed orderability’ (BDO) index that is a real number bounded between -1 and 1 (Methods). If a neuron *i* is consistently active earlier than a second neuron *j* in all trials, the neuron pair [*i,j*] is assigned a BDO of 1 (or -1 for [*j,i*] as BDO is antisymmetric) (**Fig. 7d**). Likewise, if the first neuron is active earlier than the second neuron in half of the trials but later in the other half, this pair is not orderable and is assigned a BDO value of zero. We computed BDO for all pairs of neurons, and sorted neurons by their average BDO (**Fig. 7e** for example mouse; for all mice see **Supplementary Fig. 12**). To quantify whether the overall orderability observed exceeded that of shuffled data, we computed absolute value BDO averaged over all neuron pairs (ABDO), further averaged it over all mice, and compared it to shuffled data. We found that the average ABDO value across all neuronal pairs was higher compared to that of shuffled data in all periods of the task (p < 0.001), including the two phases of the maintenance period. (**Fig. 7f**). We also found that these orderability patterns can be fully explained by temporal preference of individual neurons to earlier or later parts of a given phase, as opposed to a more complex hypothesis of time-invariant orderability, where individual cells would be orderable without having temporal preference (data not shown).Taken together, our results show that mPFC→dmStr projection neurons display orderable activity patterns in all task phases, including the delay period, and thus indicate the presence of neuronal sequences in this specific population not only during maze runs but also during WM maintenance.

Finally, we tested whether the activity of mPFC→dmStr projection neurons display significant encoding of either choice behaviour (left vs. right turn in choice runs) or performance outcome (correct vs. mistake; considering left and right sample runs separately; Methods). Across task periods, we found a relatively high fraction of neurons predictive for left vs. right turning in the encoding and retrieval periods but not in the maintenance period (**Supplementary Figs. 13**). This result corroborates the idea that for left vs. right turns partially distinct subpopulations are active (**Fig. 6e** and **Supplementary Fig. 8**). With respect to performance outcome, we found a high fraction of neurons predictive for outcome (correct vs. mistake) only during the late retrieval period (**Supplementary Figs 13**). These findings are consistent with the notion that mPFC→dmStr projection neurons participate in maintaining the relevant information for spatial working memory.

## DISCUSSION

Our study contributes to the long-standing goal of dissecting the brain circuits involved in different aspects of working memory. Whereas several key pathways have been identified and characterized, other pieces of the puzzle are still missing. For example, neurons in both dorsal and ventral hippocampus (HPC) interact with mPFC neurons in spatial WM tasks, with CA1 activity typically leading mPFC firing^35,36^. Hippocampal–prefrontal afferents engage particularly in the encoding phase of spatial WM and not during maintenance or retrieval^7^. The nucleus reuniens of the thalamus (RE) coordinates the coherent interactions across the mPFC–RE-HPC circuit^37,38^. Projections from mediodorsal thalamus (MD) to mPFC sustain prefrontal activity during the maintenance phase^8,39^. The reciprocal mPFC→MD pathway, in contrast, mainly supports subsequent choice and action selection^8,40^. Together, these results suggest that MD reinforces activity in mPFC neurons that encode task-relevant information essential for adaptive behaviour^8,40^.

Here, we complement the emerging picture of brain circuitry for WM by addressing downstream effects and specifically demonstrating that mPFC→dmStr projection neurons engage during WM maintenance. Optogenetic inhibition of mPFC→dmStr activity reduced task performance specifically when applied during the maintenance period. Using two optical population recording techniques— without and with cellular resolution—we revealed that the mPFC→dmStr pathway shows higher activity during the late maintenance period in correct trials compared to mistake trials. Despite temporally sparse activity during the maintenance period and high trial-to-trial variability, neurons displayed significant temporal orderability and showed consistent preference for specific task phases in correct trials, leading to a sequence-like temporal activation pattern. During mistake trials, this temporal order of maintenance neurons at least partially fell apart. In agreement with this finding, mPFC→dmStr bulk signals—and to some degree the population activity patterns resolved at cellular resolution—encoded information about upcoming choice in the late maintenance period (correct vs. mistake choice). Such encoding was mainly evident for upcoming choice turns to the side contralateral to the recording site. In agreement with a previous report^8^ we found weak evidence for spatial coding of mPFC neurons (right vs. left correct choice) during the maintenance period, whereas it was clearly present for encoding and retrieval periods. In contrast, other studies previously reported that a fraction of delay-related mPFC neurons is tuned to spatial information^16^ and that the direction of the sample run could be decoded from the delay-period activity of mPFC neurons^15^. In yet another study, half of the recorded neurons in dmStr significantly encoded the sample stimulus at some point during the delay period^14^. Overall, the neural basis of how space and choice are encoded in mPFC microcircuits in alternation tasks remains unclear. Resolving this issue will require a more fine-grained mapping of mPFC areas and better dissection of specific subpopulations and their interactions with other brain regions.

Nevertheless, our findings suggest that mPFC neurons functionally organize downstream striatal activity throughout the entire WM maintenance period. Sustained mPFC activity during the maintenance period may create a ramping-up activity profile in dmStr, bridging the phase of short-term memory encoding to the subsequent action (choice) similar to the situation described for evidence accumulation tasks^41^. This notion is in line with previous research demonstrating that persistent input from mPFC to dmStr represents decision variables^42^ and controls temporal processing in dmStr^43^. Because neural sequences also have been found in striatum for tasks not involving spatial WM^44^ and because sequential delay activity in dmStr was dissociated from stimulus encoding activity^14^, delay-period spanning neural sequences in dmStr may have a more general role in preparatory activity for future choices rather than serving solely working memory.

Striatal activity correlates with successful execution of goal-directed actions^24^ and our data support the idea that mPFC→dmStr activity contributes to correct action initiation following the WM maintenance period. Growing evidence indicates that mPFC and dmStr jointly contribute to successful execution of WM tasks^13^. For example, optogenetic activation of adenosine A_2A_ receptors in dmStr selectively impairs WM performance when applied during maintenance and retrieval but not during encoding^9^. Clearly, dmStr is not the only downstream region that receives information from mPFC during WM maintenance. Other potential target regions include VTA^45^ and the posterior parietal cortex (PPC)^46,47^. The specific roles of these pathways during the distinct WM periods warrant further investigation. A full picture of the dynamic interactions within the hippocampal-prefrontal-striatal-thalamic loops and beyond has yet to be established. Our newly devised multi-fiber photometry approach, which enables simultaneous photometric recordings from several tens of brain regions^48^, could be particularly well suited to measure large-scale signal flow during WM phases across multiple regions interacting with mPFC. Multi-regional recordings could elucidate how coordinated activity in distributed circuits involving mPFC and multiple striatal regions may in general govern reward-related learning of goal-directed behaviours^49^.

In addition to planning choice-related actions, prefrontal-striatal projections could be involved in impulsivity control, which is crucial for the successful execution of a WM task. In line with this notion, mPFC→dmStr projection neurons recently were found to selectively engage in inhibitory control^50^. In our experiments, perturbation of brain dynamics with the NMDA receptor blocker MK-801 induced hyperactivity, possibly by affecting such inhibitory control function of the mPFC→dmStr pathway. Consistently, pharmacologically induced impulsive behaviour was partially restored and WM performance partially rescued by optogenetic enhancement of the mPFC→dmStr pathway. Indeed, MK-801 application was previously shown to increase decorrelated activity in mPFC neurons and to decrease organized burst activity induced by coordinated synaptic inputs^51,52^. We hypothesize that the extra 20-Hz phasic mPFC→dmStr pathway activation may have helped to restore organized cortical input to the downstream striatal circuits and thereby promoted correct action choices.

More specifically, we investigated the modulatory function of HCN1 channels, which are highly expressed in mPFC and hippocampus^28^. WM impairment in schizophrenia patients has been associated with the dysregulation of HCN channels, which are therefore increasingly recognized as an important therapeutic target for controlling WM dysfunction in neuropsychiatric disorders^53^. Complementing previous research^29,30^ we demonstrated that blockade of HCN channels enhanced WM-related activity in mPFC→dmStr projection neurons. However, we did not observe improved performance in the spatial WM task, which may be due to inhibitory regulatory mechanisms or the already high performance level of healthy mice. We reasoned that a behavioural effect might become obvious under conditions of WM impairment. Application of MK-801 induces a robust WM impairment^20,21^ and can be considered to provide a relevant schizophrenia symptom animal model by blocking NMDA/glutamatergic signalling^22^. Indeed, co-administration of the HCN-channel blocker J&J12e partially rescued WM impairment induced by MK-801. A possible mechanism for this improvement could be an increased glutamate release probability, enhancing AMPA receptor-mediated, but not NMDA receptor-mediated, synaptic transmission, as was found during the co-application of the HCN-channel blocker ZD7288 and MK-801 on hippocampal slices^54^. Our results further motivate the use of HCN-channel blockers to alleviate disease conditions that impair WM. Further studies are necessary to better understand the synaptic and circuit *in vivo* mechanisms of how these blockers affect working memory functions. Focusing on the function of HCN channels specifically in prefrontal-striatal projection neurons might be a promising avenue.

## Acknowledgments

We thank Stefan Giger, Hansjörg Kasper and Martin Wieckhorst for technical assistance, Lazar Sumanovski for assistance in brain histology, Ariel Gilad and Peter Rupprecht for advice on data analysis. We thank Yasir Gallero-Salas for helpful discussions and Philipp Bethge for comments on the manuscript. This work was supported by a Transfer Project and an IPhD Project from SystemsX.ch (51TP-0_145729 and 51PHP0_157359; to F.H.), and through a Roche Joint Collaborative Project. Further support came from the European Research Council (ERC Advanced Grant BRAINCOMPATH, project 670757 to F.H.), the Swiss National Science Foundation (310030_192617 to F.H.; CRSII5-173721 and 315230_189251 to B.F.G.), the German-Swiss Research Unit “Barrel Cortex Function” (DFG FOR1341, SNSF 310030E-147485; to F.H.), the Human Frontiers Science Program (RGY0072/2019 to B.F.G), the ETH Zurich (ETH-2019-01 to B.F.G.), and the University of Zurich (Forschungskredit to C.L.).

## Author Contributions

M.C. and F.H. conceived the project and designed all experiments. J.L.A.W. automated the T-maze alternation task. M.C. carried out all experiments except multi-unit electrophysiology recordings which were performed and analysed by C.L.. L.S.C. assisted in fiber-optic related surgeries and experiments. M.C., Y.S., and F.H. analysed the fiber photometry data. R.B. and A.E. performed the surgeries and assisted in the miniscope experiments. A.F. contributed to photometry analysis and performed the majority of the analysis for miniscope experiments. B.H., E.O., Y.S., A.F., and B.G., assisted in the study design and interpretation of experimental results. M.C., Y.S. and F.H. wrote the paper with comments from all authors.

## METHODS

### Animals

Experiments were performed on 34 male C57BL/6 mice and 4 male transgenic mice Ai148(TIT2L-GC6f-ICL-tTA2)-D (Nr. 030328, The Jackson Laboratory) for the miniscope experiments, all aged 6–8 weeks at the first use. Mice with chronic implants were housed individually under a reversed 12-h L/light–dark cycle with food and water available *ad libitum* before behavioral training. During behavioral training, mice were water-restricted and maintained at 85% of their initial body weight. All experiments were performed during the animals’ dark period. Experimental procedures were conducted in accordance with the guidelines from the Veterinary Office of Switzerland and were approved by the Zurich Cantonal Veterinary Office.

### Surgical procedures

Mice were first anesthetized with isoflurane (1.5 – 2%) and head-fixed in a stereotactic apparatus. The viral injections and fiber implantations were performed in the same surgery. First, after the small craniotomy, retrograde virus AAV-retro/2-hSyn1-chI-mCherry_2A_NLS_iCre (210 nl, ∼1×10^9^ vg μl^−1^) was delivered unilaterally into the left dmStr (+1.05 AP, -1.50 ML, -2.05 DV), corresponding to the medial anterior part of dmStr^10^. On the same day, another viral injection was delivered into the left PrL (+1.8 AP,-0.6 ML, -1.3 DV): we used AAV2.9.Syn.Flex.GCaMP6m (210 nl, ∼1×10^9^vg μl^−1^) for calcium recording experiments, AAV2.5-CAG-FLEX-ArchT-GFP (210 nl, ∼1×10^9^ vg μl^−1^) for inhibition experiments, and AAV2.2-EF1a-DIO-hChR2(E123T/T159C)- mCherry (210 nl, ∼1×10^9^ vg μl^−1^) for activation experiments. For fiber photometry and optogenetic experiments, mice were unilaterally implanted with a ferrule-coupled optical fiber (0.22 NA, 400-μm diameter) after viral injections in the same surgical session. The distal end of the inserted fiber implant was cut and polished at a 45° angle and the flat side positioned immediately dorsal to the PrL area and directed towards the brain’s midline. Finally, the fiber implant was fixed in place by glueing it to the skull with dental cement. For the miniscope experiments we performed a second surgery 12-14 days after the virus injection. For insertion of the GRIN lens (Gradient Index Lens; GRINTECH, Jena, NEM-100-06-08-520-S-0.5p; 1-mm diameter, 80-μm working distance, 4.54-mm length), we implanted a custom-made stainless steel guide cannula (1.2-mm outer diameter) with a glass coverslip (0.125-mm thick BK7 glass, Electron Microscopy Science) glued to the bottom. After cleaning the skull from periosteum and tissue, we performed a circular craniotomy of 1.3-mm diameter, centered above the mPFC (1.9 mm anterior and 0.4 mm lateral to Bregma,). We aspirated 1.2 mm of brain tissue overlying the targeted PrL area to prevent increased intracranial pressure when inserting the cannula. The guide cannula was lowered 2.3 mm (from the skull surface) into the craniotomy and fixed with UV-curable glue (Loctite 4305). For secure attachment and durability of the implant, two anchor screws (18-8 S/S, Component Supply) were placed on the contralateral parietal and the interparietal plate. Next, either blue-light curable Scotchbond Universal (3M ESPE) or Metabond (Parkell) were applied around the implant, the screws, and the exposed cranium. For additional stability and for attaching a small metal head bar to the implant, a layer of dental acrylic (Paladur) was added on top of the first layer. Mice received postsurgical analgesic and anti-inflammatory treatment for three days with buprenorphine (0.1 mg/kg BW, s.c. every six hours during daytime, and 0.01 mg/ml in drinking water during the night) and carprofen (4 mg/kg s.c., twice a day).

### Histology

At the end of each experiment, mice were transcardially perfused with 4% paraformaldehyde in phosphate buffer, pH 7.4. Fixed tissue was then sectioned (100 μM) using a vibratome and mounted on slides with Fluoromount (Dako). Direct fluorescence of GCaMP6m, GFP-tag of ArchT or mCherry tag of ChR was then examined under a confocal microscope (Fluoview 1000; Olympus) to assess the extent of viral spread and the axon-terminal expression pattern.

### Behavioral setup and training

Behavioural experiments were performed using a data acquisition interface (USB-6008, National Instruments) and custom-written MATLAB software to control devices required for the task and to record trial-related neural activity and licking data. Behaviour training for the recording experiments started ∼2 weeks after the surgery (∼5 weeks for inhibition and miniscope experiments). First, mice were given 2 days of habituation to the T-maze. Habituation sessions consisted of 5–10 min of free exploration in the maze with all doors open and freely accessible water rewards. On the subsequent 2-5 days, mice underwent behavioural shaping, which consisted of 10 trials per day running to baited goal arms in alternating directions. A drop of water (10 μl) was used as a reward. Reward delivery was triggered by licking and controlled with a miniature rocker solenoid valve (0127; Buerkert). For the miniscope experiments, mice were also accustomed to carry the miniscope by habituating them to its weight in 20-min habituation sessions in the home cage. Once mice followed the sequence of the shaping procedure at a speed of 1 min per trial, they underwent further training in the T-maze task until they reached the desired performance criterion (>60% correct trials on 2 consecutive days). In the training phase, mice in the choice run had to choose the T-maze arm (right or left) opposite to the arm they had visited in the sample run before the delay. Each trial of the session consisted of three periods: encoding, maintenance and retrieval. In the encoding period (sample run) one of two goal-arms was blocked by a door and the mouse was directed towards a water reward in the open arm. During this period, the animal must encode the location of the sample reward. In training and recording sessions, sample arms were pseudorandomized (not more than three consecutive runs to the same sample arm). In the maintenance period, mice returned to the start box and had to maintain the sample reward’s location in their working memory for a delay period (5-s duration for fiber photometry and miniscope experiments; 10-s duration for optical and pharmacological perturbations). In the retrieval period (choice run), all doors were opened and the mouse had to select the previously closed goal-arm to receive a second reward. Inter-trial delay was always fixed at 10 s. Performance of mice was maintained under 80% by providing 2-3 training sessions per week.

### Pharmacological interventions

First, mice were trained in the T-maze alternation task to reach the desired performance criterion: 2 days >60%. Next, mice were accustomed to the stress of systemic i.p. injection using the vehicle (0.3% Tween 80 in 0.9% NaCl; 10 ml/kg injected volume) 30 min before the beginning of the experiment. When mice reached the performance criterion under the vehicle injection conditions, they underwent the test day experiments with i.p. injection of either MK- 801 (0.1 mg/kg) or J&J12e (3 mg/kg) or both drugs given in combination, 30 min prior to the test experiment. A fresh compound solution was prepared on each test day in the vehicle solution (0.3% Tween 80 in 0.9% NaCl; 10 ml/kg injected volume), sonicated for 30 min, and well mixed. In the first experimental design (Fig. 4b, Supplementary Figure 4a,b), we trained mice in the alternation task with a 5-s delay, followed by the baseline day with the same 5-s delay (after vehicle injection). Next, we tested the mice’s performance with a challenging 10-s delay after systemic injection of either the J&J12e compound or vehicle (J&J12e or Vehicle probe days respectively). To balance cohorts of mice, 50% of mice underwent vehicle testing first, followed by testing with J&J12e, and the other 50% of mice started with J&J12e testing, followed by vehicle testing. In the second experimental design (Fig. 4c), the delay was kept at 10 s for all test days. To balance cohorts of mice, 50% of mice underwent the MK-801 test first, followed by testing with the J&J12e and MK-801 in combination; for the other 50% of mice, the test order was reversed. Following the first test day, mice were given a 5-6 days break in their home cage without experiments.

### Fiber-photometry and optogenetic manipulation

Pathway-specific photometric recordings were carried out through the implanted 400-μm diameter optical fiber using 0.3 mW of 488-nm excitation light (OBIS LX 488-nm laser, Coherent), modulated at 490 Hz. The 425-nm light (LuxX 425 laser, Omicron) used to control for motion artefacts was modulated at 573 Hz and delivered together with the 488-nm laser light through the same fiber. The dichroic beam-splitter (No. F58-486, AHF) directed excitation light into the objective (F240FC-532, Thorlabs) and transmitted fluorescence signals. The fluorescence from the GCaMP6 expressing neurons was propagating along the optical fiber towards the photometry setup, was collimated by the same objective (F240FC-532), passed the emission filter (525/50 nm, No. F37-516, AHF) and was focused by the condenser lens (ACL3026U-A, Thorlabs) onto a PMT (H10770(P)A-40, Hamamatsu). The photocurrent was amplified in the custom built pre-amplifier and transmitted to the data acquisition board (DAQ USB 6211, National Instruments). The fluorescence signals were recorded at 2 kHz continuously until the end of each session and demodulated with the digital lock-in detection (MATLAB R2019b). Fluorescence contributions excited by the 425-nm and 488-nm lasers were demixed by demodulation at the respective modulation frequencies in 50-ms time bins, resulting in an effective 20-Hz rate for the fluorescence recording. At the beginning of each recording session, the mouse was placed into the start box before the first trial. Mice were simultaneously video tracked with the IR illuminated camera (DMK 33UX178, Imaging Source). Behavioral triggers (*e*.*g*., for opening/closing maze doors and triggering the water valve to open) were set and recorded simultaneously with the fluorescence signals when mice entered regions of interest in the maze (ROIs). Therefore, we could online time-stamp our recording to all phases of the T-maze alternation task. All continuous behavioural parameters such as the licking response were also simultaneously recorded on the same data acquisition board (DAQ USB 6211, National Instruments). All behavioural analysis was performed using custom-written MATLAB scripts. First, based on the behaviour videos and licking actions, we identified 11 salient events in each of the 1085 trials collected from 6 mice. For correct trials the events were the following (Fig. 1a): 1–start of the sample run, 2–turning at the T-junction of the main maze arm, 3–first water reward, 4– end of licking period, 5–turn to run towards the start box, 6–reaching the start box, 7–start of the choice run, 8–turning at the T-junction, 9– second water reward, 10–end of licking period, 11–end of choice run. Because mice did not receive a reward in incorrect trials, events 9 and 10 were omitted for these trials.

Pathway-specific optogenetic inhibition experiments were carried out using constant illumination with 20-mW, 561-nm light for activation of ArchT. For pathway-specific activation in ChR2-expressing mice we applied 20-Hz pulses of 10-mW, 473-nm light, delivered through a 400-μm diameter optical fiber (NA 0.22). In each session employing optogenetics, perturbation light was delivered in 50% of randomly interleaved trials and targeted exclusively at one of the task periods (e.g., only during encoding period for one session and only during maintenance in a different session). In total, each mouse experienced 9 sessions, with 3 perturbation sessions for each task period.

### In vivo electrophysiological recordings

We validated optogenetic excitation and silencing of mPFC→dmStr projection neurons by performing acute in vivo electrophysiology. Mice (n = 4) were anesthetized with isoflurane (2% for induction; 0.8% during recording) and body temperature was maintained at 37°C using a heating pad. A small craniotomy (1-mm diameter) was performed to provide access to the left PrL and the brain was covered with silicon oil. A silver wire was placed in contact with the cerebrospinal fluid through a small (0.5 mm) trepanation over the cerebellum to serve as a reference electrode. A silicon probe (Atlas Neurotechnologies, 16 linear sites, 100-μm spacing) was inserted through the craniotomy into the left cortical hemisphere to record multi-unit activity from the injection site in the left PrL and surrounding cortex. After insertion of the silicon probe, we positioned an optical fiber (0.2-mm diameter) directly next to the probe, directed downwards towards the brain. We waited 30 minutes to allow the recording to stabilise after implantation of the electrode array. After stabilisation, the broadband voltage was amplified and digitally sampled at a rate of 30 kHz using a commercial extracellular recording system (RHD2000, Intan Technologies). The raw voltage traces were filtered offline to separate the multi-unit activity (MUA; bandpass filter 0.46-6 kHz) using a fourth-order Butterworth filter. Subsequently, the high-pass data were thresholded at 5.5 times the standard deviation across the recording session and the numbers of spikes in windows of interest were counted. Once the recording was stable, we performed light stimulation through the optical fiber to drive the expressed opsin. We delivered light by connecting the fiber-optic cannula via a fiber-optic patch cable to a laser (either OBIS 473 nm LX for ChR2, or OBIS 561 nm LS for ArchT). The CW laser was gated in a stepwise manner with a function generator, triggered by a TTL-pulse from the electrophysiology computer. We found cells directly driven by the laser stimulation, as well as cells presumably showing secondary effects via connectivity with driven cells. To combine data across mice, MUA activity at sites with clear light-induced responses was expressed as a percent change from the baseline, i.e. the average spike rate during the 10-s pre-laser stimulation period (100%). All MUA activities were combined from the injected or control region and a t-test was performed between the baseline period (10-s before stimulation) and the stimulation period (during laser onset).

### Photometry analysis

Calcium indicator fluorescence signals were expressed as percentage change in fluorescence ΔF/F=(F(t)- F_0_)/F_0_, where F(t) denotes the fluorescence value at time t across the entire trial time (from event 1 to event 11) and F_0_ the baseline fluorescence level calculated as the mean value in the 1-s time window before the maintenance period (before event 5). For analysis across trials and mice, we z-scored the fluorescence signals for individual trials by calculating (F(t)-F_0_)/σ, where σ is the standard deviation of the fluorescence values in the 1-s baseline window for F_0_ calculation. Because mice behaved freely in the T-maze, trial phases varied in their duration. To temporally align trial-related signals we therefore resampled the z-scored signals for each trial phase to match the median phase duration across all trials. This normalization by temporal resampling permitted us to average and compare trial-related activity across trials and mice. All mice contributed 4 expert sessions to the dataset, which in total comprised 1085 trials (544 rightward sample runs: 407 correct, 137 mistakes; 541 leftward sample runs: 405 correct, 136 mistake). The resampled traces were used for detailed analysis of the maintenance period (between events 5 and 7) in Figures 2 and 4.

### Miniscope imaging and data analysis

Four weeks after GRIN lens implantations, mice with GCaMP expression were taken into behavioural experiments. We used a head-mounted miniaturized microscope (nVista, Inscopix) for calcium imaging with cellular resolution. All mice were recorded at 20 Hz with a gain of 3 to 4 and 10-25% LED illumination (0.2 to 0.5 mW/mm^2^). Simultaneously, we video tracked the mouse and recorded behavioural parameters such as water reward delivery and licking response. After mice completed several expert sessions, calcium imaging data were pre-processed using Inscopix Data Processing Software (version 1.2.1), allowing for motion correction and extraction of cellular calcium signal traces. For efficient data processing, movies were spatially downsampled by a factor of 4 and temporally downsampled by a factor of 2. After applying motion correction in the Inscopix software, ΔF/F was calculated pixelwise using the mean fluorescence across all frames as F_0_. Then, we applied the Inscopix PCA-ICA algorithm to automatically identify the spatial location of cells and their activity profile throughout the recorded session. Finally, in order to match cells recorded over several sessions, we employed the longitudinal registration algorithm. For further analysis, we used only those cells that were successfully recorded longitudinally in all expert sessions. For comparison with the fiber-photometry data, we summated the pre-processed activity of all longitudinally tracked neurons (28 – 88 neurons per mouse) for each mouse (n = 6 mice, 3-5 sessions per mouse) and treated these population signals in the same way as the bulk fluorescence signals measured by fiber photometry.

### Analysis of single-cell activity across task periods and task phases

We evaluated the degree to which neurons were specifically active in particular time windows (either the 3 task periods or the 10 task phases defined as the intervals between the 11 salient T-maze events). To optimize temporal accuracy and reduce the amount of non-spike fluctuations in the signal, we temporally deconvolved the pre-processed miniscope calcium imaging data using a supervised algorithm based on deep neural networks^34^. Next, we calculated for all neurons and each trial the mean activity in each time window considered (either periods or phases). Based on a rank sum test of mean activity values for each time window across all correct trials against the pool of all other time windows, we identified the significance level of a given neuron being active in a particular time window. The significance level is presented as –log_10_(p), where p is the p-value of the rank sum test (e.g. if p=0.01, –log_10_(p) = 2). To quantify the fraction of neurons significantly active in specific time windows (or pairs of time windows), we binarized the significance levels by thresholding at p = 0.01. For either task periods or task phases, we plotted these neuronal fractions as a symmetric matrix, with values on the diagonal representing the fraction of neurons displaying significant activity in at least this time window, and off-diagonals showing the fractions of neurons active in both of the respective time windows.

### Neuronal encoding of behavior

To analyse encoding of behaviour by the bulk activity of mPFC→dmStr projection neurons (based either on the photometry data or the summated cell activities for miniscope imaging) we used receiver operating characteristics (ROC) analysis. We considered three behavioural aspects: left vs. right turning direction and correct vs. mistake performance in choice runs, considered separately for left and right sample runs. We used the z-scored calcium signals for all expert sessions (>60% correct trials on 2 consecutive days), resampled to align them to the task phases. We also applied ROC analysis to the experimental data with pharmacological intervention (J&J12e correct vs. mistake; and J&J12e vs. Vehicle; Fig 4 and Supplementary Fig. 4). For each pair of trial types considered, we randomly selected 100 trials for each group (20 times repeated) and calculated the area-under-the-curve (AUC) for the ROC curve. We performed this analysis for each time bin across the full time window of interest using a 3-point moving window. To test for significance of classification of the two trial types, we additionally performed the same analysis for data with shuffled trial labels. In the figure plots we show the mean ± s.e.m. of the AUC values for real and shuffled data (for each time point and considering the 20 draws). We used a Mann–Whitney U test (p < 0.01) to evaluate the difference between real and shuffled datasets and test whether trial type classification was significant.

To evaluate neuronal encoding on the population level, based on the miniscope data, we tested for each neuron whether in a given time window (task period or task phase) its mean activity was predictive of behaviour (either for left/right turning direction or for correct/mistake performance in choice runs; ranksum test, p < 0.01). For each time window, we then counted the number of predictive neurons and tested if this number was significant by using a binomial model, in which each neuron, as null hypothesis, could independently be significant with probability p =0.01. For all mice, we plotted the fraction of cells whose activity was significantly predictive of the behavioural classification considered, e.g., left vs. right turns, for task periods and for task phases in **Supplementary Fig. 13a** and **b**, respectively.

Finally, we investigated if specific behavioural aspects were encoded transiently during the maintenance period. As for the above analysis, we focused on left vs. right turning behaviour and also analysed correct vs. mistake performance, separately for left and right sample turns. We computed the euclidean (L2) norm between neuronal population signals, using the deconvolved ΔF/F traces averaged for each of the two behavioral conditions, and tested the result against a null model with condition labels shuffled (p < 0.01, permutation test). We performed this test for each time bin of resampled trial time, and plotted the significance level as -log10(p) as a function of the resampled time (**Supplementary Fig. 14**). We also performed transient decoding using cross-validated machine learning methods: linear regression, logistic regression and 2-layer neural network (data not shown). Results were largely consistent with L2 norm for direction discrimination, but showed no effect for performance discrimination. We concluded that these methods do not converge to optimal performance due to a low amount of mistake trials.

### Orderability analysis

To quantify activation sequences in neuronal populations, we introduced the ‘Binary Directed Orderability’ (BDO) index. For each pair of neurons (*i,j*) and each trial, BDO evaluates whether most of the signal in the neuron *i* came before that of the neuron *j* or vice-versa (hence the name binary), then finds the average frequency over trials of the neuron *i* being earlier than neuron *j*. Thus, this metric determines whether neuron *i* is consistently active earlier than neuron *j* across trials, independent of the magnitude of the time differences and of the exact time period in which neurons are active. For each active neuron, we first calculated the probability *p(t)* of being active at time *t* by normalizing the baseline-subtracted deconvolved signal x(t) (after baseline subtraction) so that the sum over all time bins 1..*N* is one:

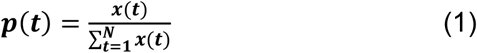

Next, we determined the center of mass *μ* of *p(t)*, i.e., the time bin in which the neuron is most active:

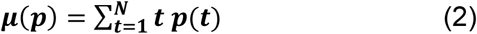

We prefer to use *μ* rather than the peak time because *μ* is more robust against small perturbations when calculated for each noisy trial, and corresponds to the weighted average time of the entire neuron trace, not of one local event. For each pair of neurons [*i,j*] and each trial *k*, we defined the binary order in (*T*_k,ij_) as one if neuron *j* was active later than neuron *i* (*µ*_j_> *µ*_i_) and as zero otherwise. Thus, the fraction of trials, in which neuron *j* was active later than neuron *i*, is given by

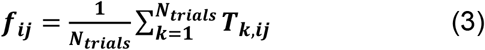

where *N*_*trials*_ is the total number of trials. Because the case *µ*_j_ = *µ*_i_ can be neglected for floating point comparison, it follows that *f*_ij_ + *f*_ji_ = 1.

Finally, the BDO index is defined as the linear function of the above fraction

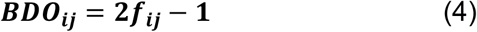

The BDO index is bound between [-1, 1]. It is zero when no consistent order occurs across trials, +1 if neuron *j* consistently follows neuron *i*, and -1 if neuron *j* leads neuron *i*. For visualization, we plotted the BDO for all neuron pairs as a heatmap, where the hue denotes BDO value. For validation, we compared this plot to a shuffled heatmap, for which a shuffle over neuron labels was performed for each trial individually. To quantify whether BDO was above chance, we defined the average absolute value BDO (ABDO) as the mean absolute value of the off-diagonal BDO values

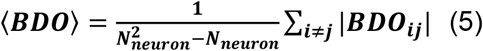

where *N*_*neuron*_ denotes the total number of neurons. If <BDO> is significantly larger than for the shuffled heatmap (p ≤ 0.01, permutation test), this means that the signals recorded from neurons in this population are on average better pairwise orderable than random signals.

## SUPPLEMENTARY FIGURES

**Supplementary Figure 1.**
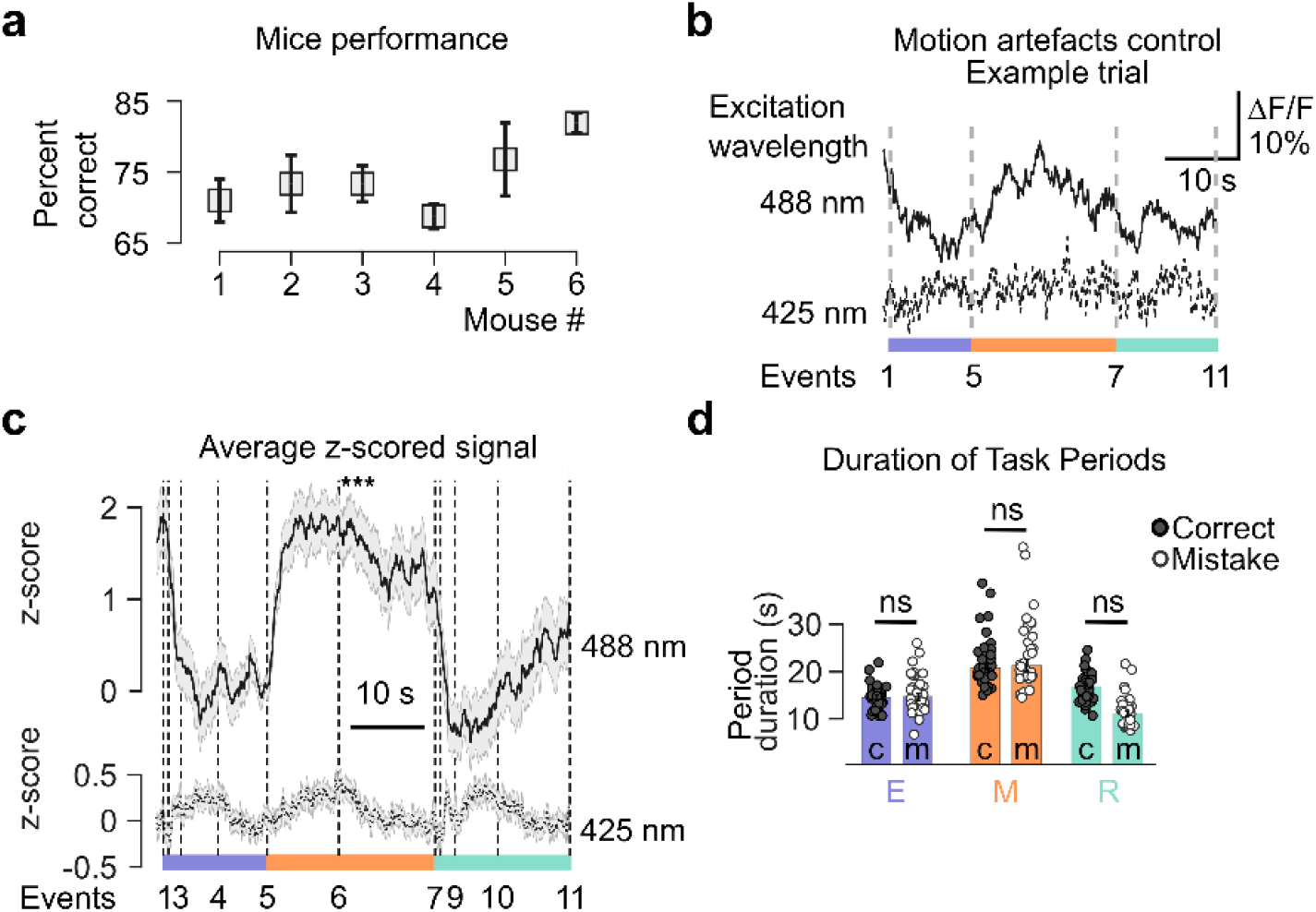
Behavioral parameters and motion-artefact control for photometry experiments in mice performing T-maze alternation. (**a**) Expert performance levels of the mice used in the fiber photometry experiments shown in Figs. 1 and 2. (**b**) Single-trial example ΔF/F trace with 488nm excitation (upper trace) and 425-nm excitation (lower trace; near-isosbestic for GCaMP6f). Traces were recorded simultaneously by delivering light through the same optical fiber and modulating each wavelength with a different frequency (573 Hz for 488-nm light and 490 Hz for 425-nm light). (**d**) Average z-scored ΔF/F signal profile for 4 mice with dual-wavelength excitation. Excitation at 425 nm produced traces with significantly lower variance compared to 488-nm excitation, confirming the calcium-dependence of the observed signals and excluding major contributions of intrinsic signal changes or motion artefacts. The mean z-score over the maintenance period was significantly higher for 488-nm compared to 425-nm excitation (***, p < 0.001, paired Wilcoxon signed-rank test). (**d**) Duration of the three salient task periods (marked by purple, orange, and cyan) for correct trials (black dots) and mistake trials (empty dots). Mean durations were not significantly different (ns, p > 0.05).

**Supplementary Figure 2.**
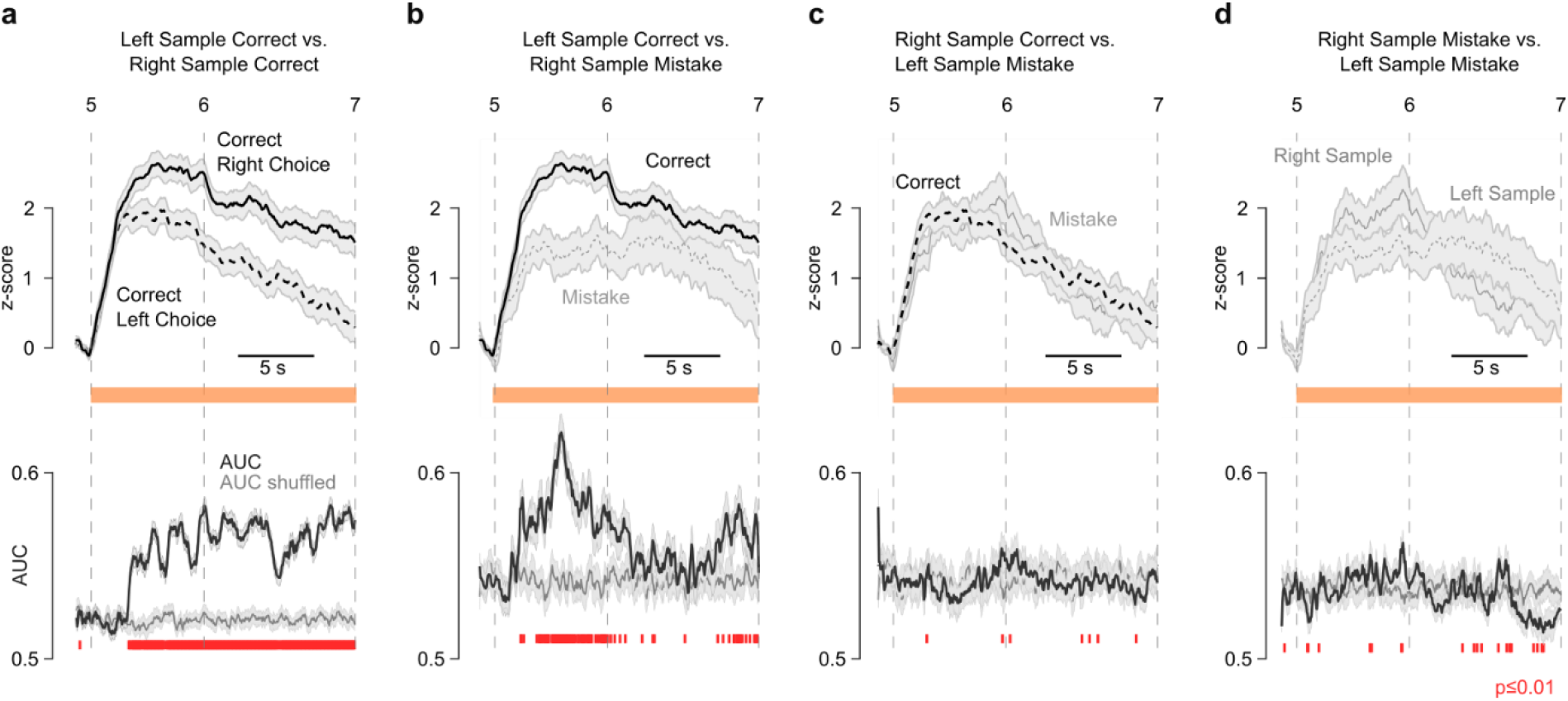
mPFC→dmStr pathway activity during WM maintenance preceding different behavioural outputs. Top row: Average z-scored ΔF/F traces of mPFC→dmStr activity during the WM maintenance period, with each plot comparing two conditions (n = 6 mice). Dashed vertical lines indicate trial events defining the maintenance period. Bottom row: Area-under-the-curve (AUC) from the ROC analysis comparing the two conditions. (**a**) Comparison of correct choices (right or left) following left and right sample runs, respectively. (**b**) Comparison of right turns following left sample runs (correct choices) and right sample runs (mistakes). (**c**) Comparison of left turns following right sample runs (correct choices) and left sample runs (mistakes). (**d**) Comparison of incorrect choices (mistakes; left or right) following left and right sample runs, respectively. For statistical analysis AUC traces were compared to AUC values for shuffled trials. Red vertical lines indicate time bins, for which a Mann-Whitney U-test shows significance at p < 0.01.

**Supplementary Figure 3.**
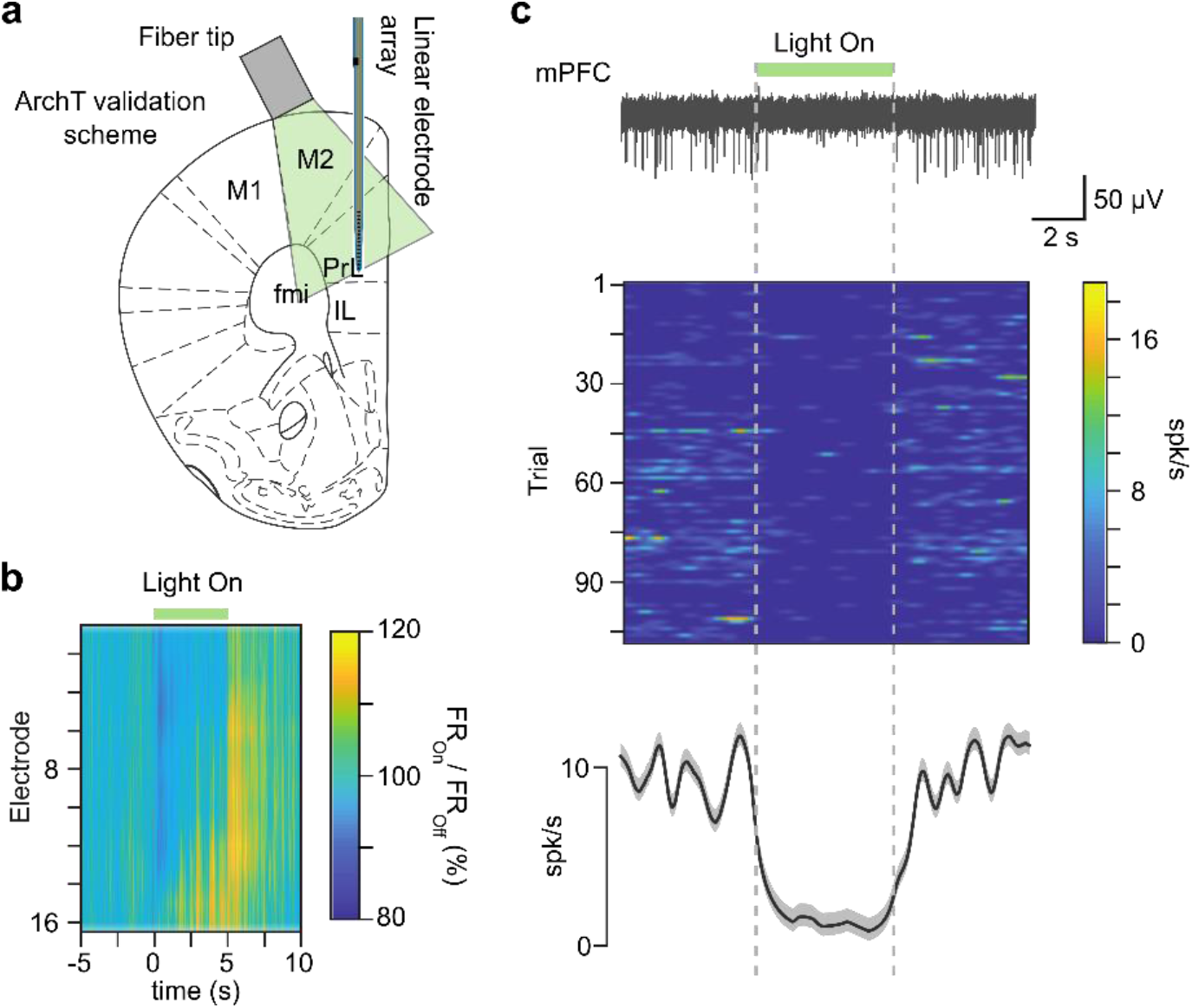
Validation of pathway-specific optogenetic inhibition using ArchT. (**a**) Schematic of experimental design: mPFC→dmStr projection neurons were infected with a viral construct expressing Cre-dependent GFP-tagged ArchT. An optical fiber was placed on the brain surface for transient inhibition of neuronal activity with 561-nm light delivery. The linear probe was inserted vertically. (**b**) Light-evoked multi-unit activity along linear array in mPFC showing spatial distribution of suppression (16 site linear array: top electrode – the most dorsal). **c**) Top: example cell in mPFC showing suppressed activity following light delivery to the surface of mPFC. Middle: average response, for n = 110 trials. Bottom: Spikes per second in a trial-wise light-modulation of the example cell.

**Supplementary Figure 4.**
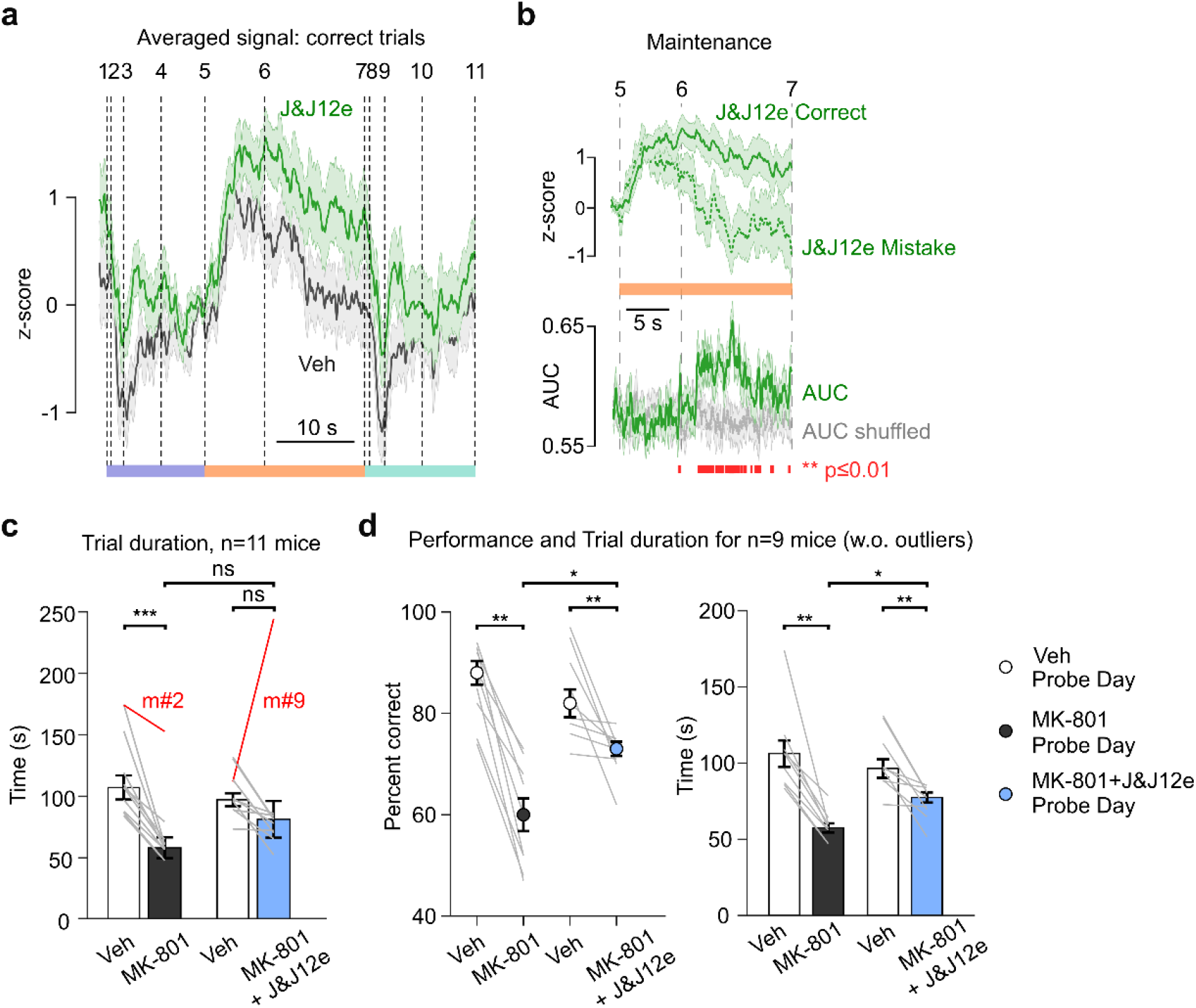
Effect of HCN channels blockade on WM-related mPFC→dmStr activity and WM impairment induced by MK-801. (**a**) Resampled z-scored calcium traces for the J&J12e condition in green and vehicle (Veh) in black (n = 6 mice). Numbers correspond to the task events indicated in Fig 1. Solid lines represent average calcium transients, the shaded areas indicate s.e.m. (**b**) Top: With J&J12e application, mPFC→dmStr pathway calcium signals during the maintenance period are higher for correct trials (solid line) compared to mistake trials (dashed line). Bottom: Time course of correct vs. mistake classification based on AUC of an ROC analysis. Red markers at the bottom indicate time bins with significant classification accuracy compared to trial-shuffled data (grey line; p < 0.01; Mann-Whitney U-test). J&J12e attenuates MK-801-induced hyperlocomotion as indicated by trial durations (***p < 0.001, paired Wilcoxon signed-rank test). Nearly all mice (9 out of 11) exhibited hyperlocomotion after receiving the MK-801. However, two mice (highlighted in red) displayed cataleptic-like behaviour (pausing in various places of the maze for several minutes), as has been reported previously (ref. 17). (**d**) Analysis of performance and trial duration for the various conditions restricted to the 9 mice that did not show cataleptic-like behaviour.

**Supplementary Figure 5.**
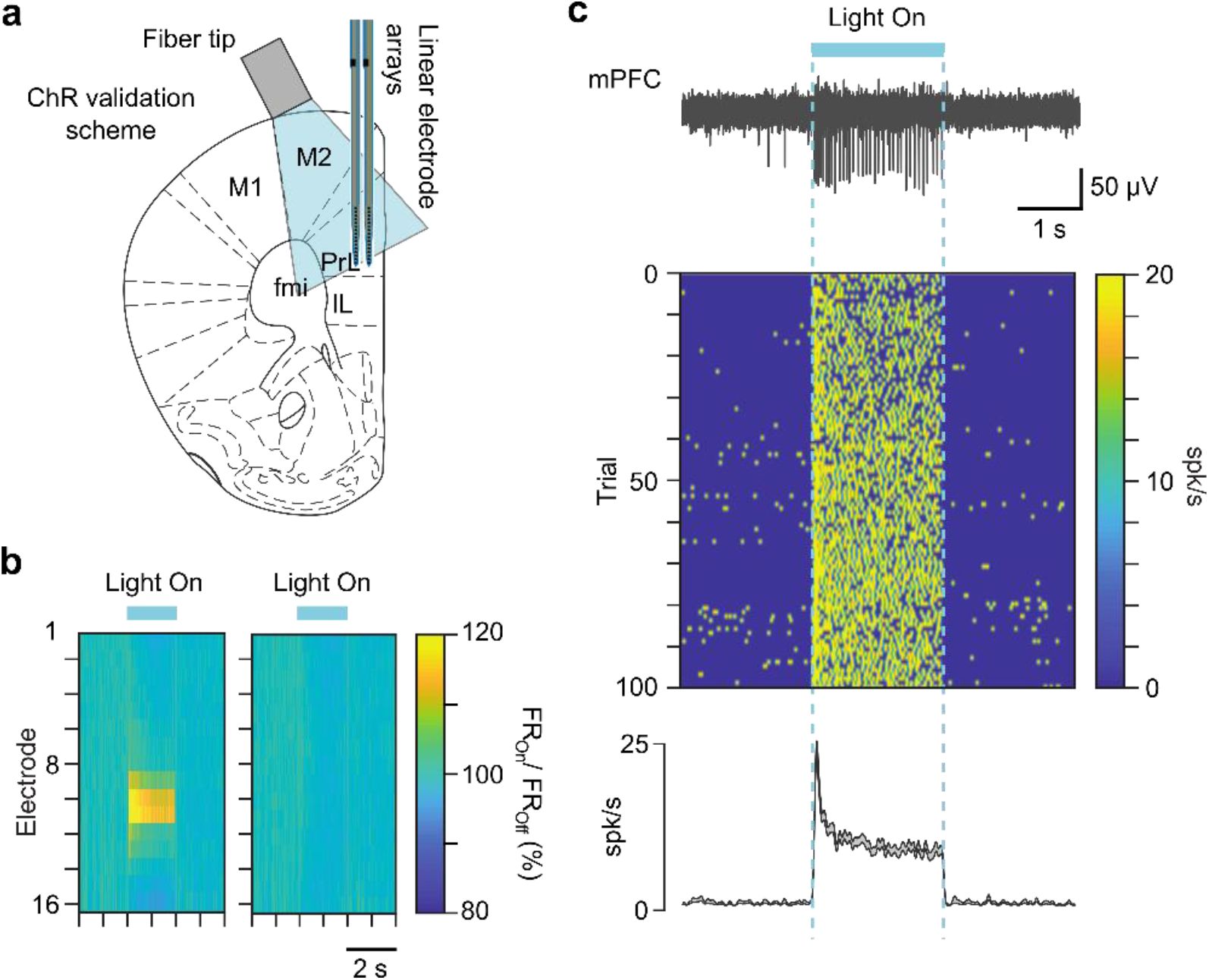
Validation of pathway-specific optogenetic activation. (**a**) Scheme of ChR2 validation experiment: similar to the Fig.5, mPFC→dmStr projecting neurons were infected by the Cre-dependent ChR2 virus injection, the optical fiber was placed on top of the brain surface for the transient activation of neuronal activity with 473 nm light delivery. (**b**) Light-evoked multi-unit activity along two linear arrays in mPFC showing spatial distribution of activation (2×16 site linear array: top electrodes are the most dorsal). (**c**) Top: Example cell in mPFC showing enhanced activity following light delivery to the surface of mPFC. Bottom: average response of the cell in (b, top) (n = 200 trials) c) Trial-wise light-modulation of the example cell (n = 200 trials).

**Supplementary Figure 6.**
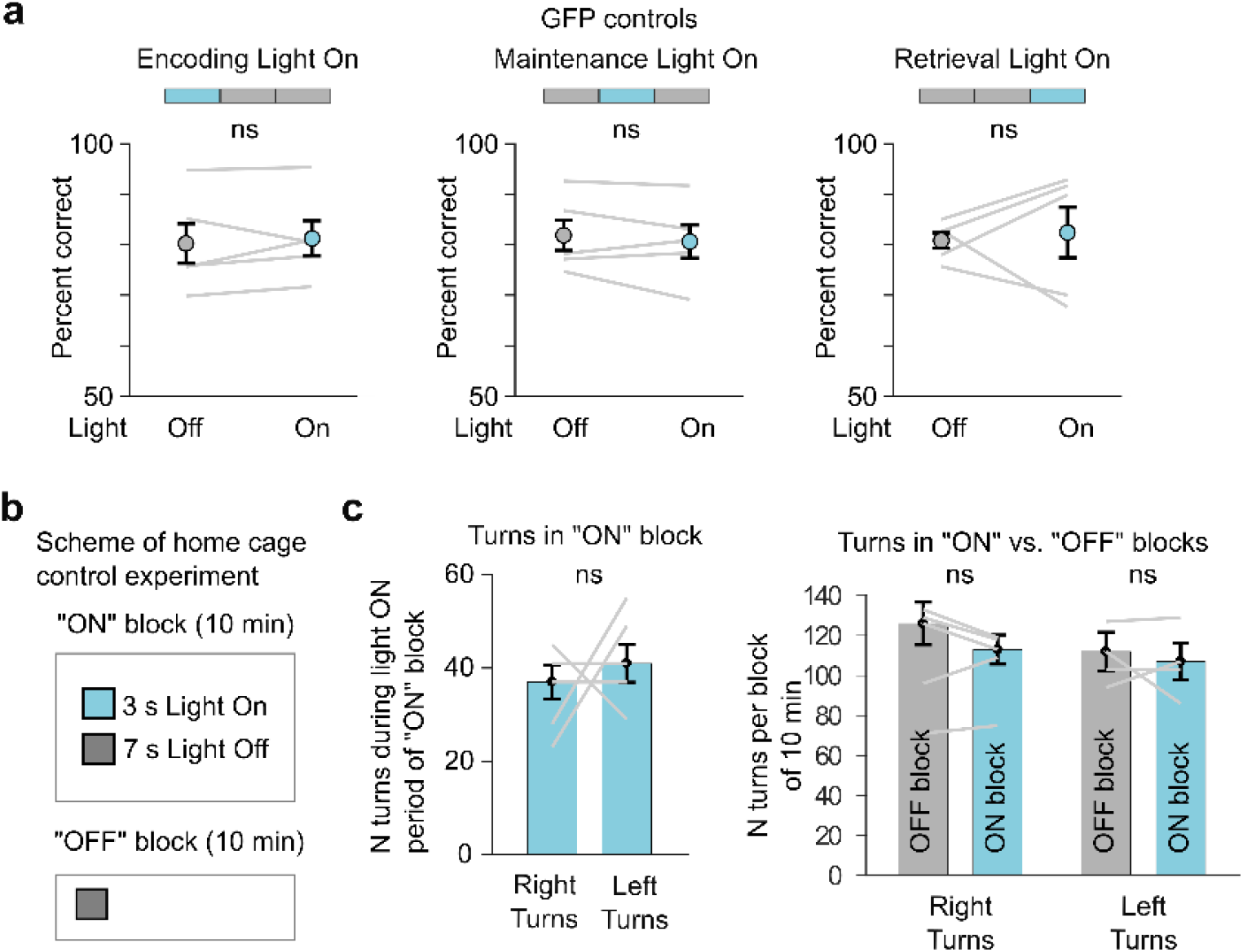
Control experiments to test for light-induced behavioural effects. (**a**) No significant effect on performance was observed upon light stimulation in control mice injected with a virus driving expression of GFP instead of ChR2 (n = 5). As for the ChR2 experiments in Figure 5, light stimulation was applied in random 50% of trials. Grey lines represent individual mice. (**b**) Design of home cage control experiments to test for potential effects of ChR2 activation on motor behaviour (n = 5 mice with ChR2 expression). Mice were tested twice in their home cage: On the first day, explorative behaviour was monitored for 10 min with continuously alternating periods of 3-s laser light on and 7-s laser light off periods (ON-block), followed by a 10-min period without no laser illumination (OFF-block). On the second day, the order of the test blocks was reversed (first 10 min OFF-block, then 10 min ON-block). Data from both days was analysed and pooled. (**c**) Left: We observed no significant difference in the number of right vs. left 90° turns inside the 3s activation period of the ON-block (*p >* 0.4, paired Wilcoxon signed-rank test). Right: The number of 90° turns within ON-vs. OFF-blocks did also not differ (*p > 0*.*4*, paired Wilcoxon signed-rank test) considering the entire 10 min duration per block; this analysis controls for potential long-lasting light-induced turning preferences that could have occurred during the 7-s light off period in the ON-block).

**Supplementary Figure 7.**
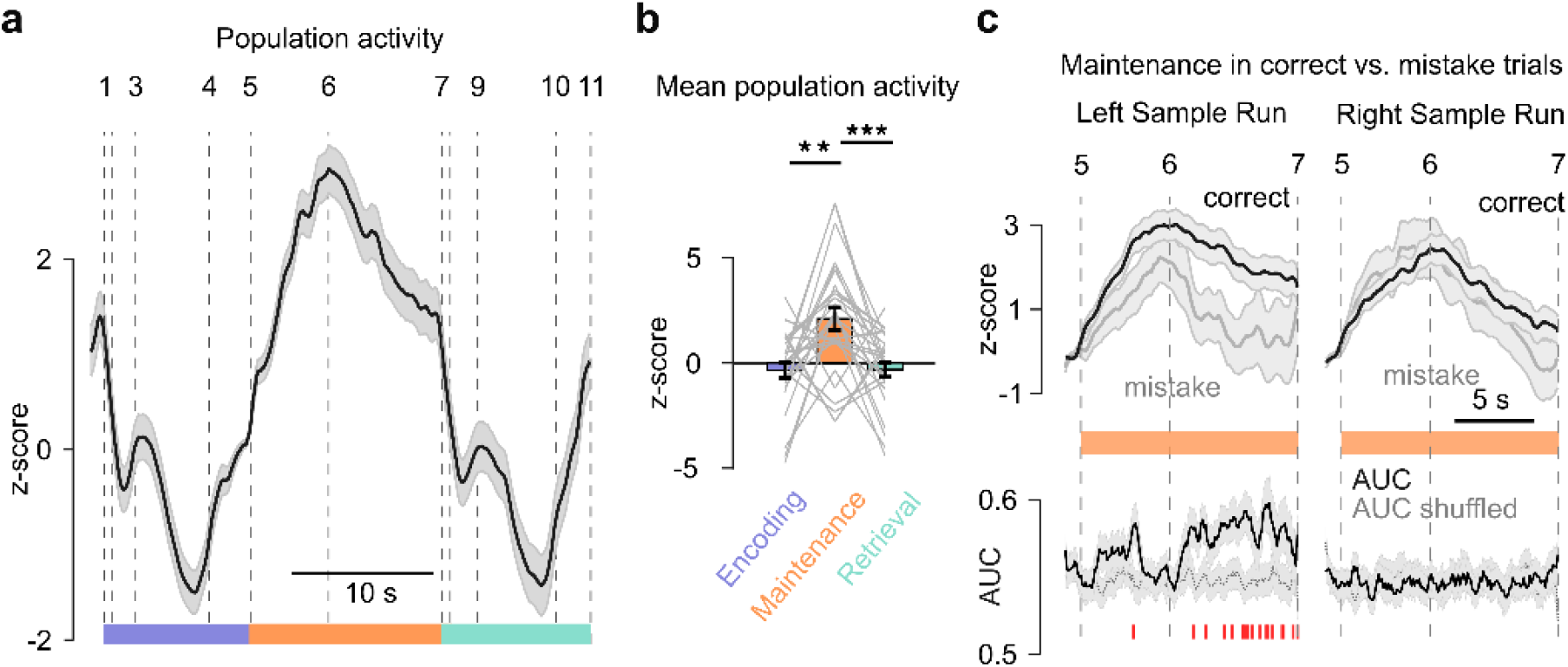
Average neuronal population activity in the miniscope imaging data. (**a**) Mean task-related calcium signal across all identified neurons imaged with the miniscope (shaded area indicates s.em.). As for the fiber photometry data, neuronal ΔF/F traces were resampled and aligned to the task phases, z-scored, and finally averaged across sessions and mice (n = 6 mice; 3-5 sessions per mouse; 28-88 active neurons per mouse were longitudinally tracked across days). Dashed lines indicate task events for alignment as defined in Figure 1. (**b**) Average activity was higher during the maintenance period compared to encoding and retrieval periods (**p < 0.01, ***p < 0.001; paired Wilcoxon signed-rank test). (**c**) Top: Mean z-scored calcium signals during the maintenance period for correct (black) and mistake (grey) trials, shown separately for periods following left sample runs (correct choice to the right, contra-lateral to the recording site) and right sample runs (correct choice to the ipsi-lateral side). Bottom: Time course of correct vs. mistake classification based on the area-under-the-curve (AUC) of an ROC analysis. Red markers at the bottom indicate time bins with significant classification accuracy compared to trial-shuffled data (grey trace; p < 0.01; Mann-Whitney U-test).

**Supplementary Figure 8.**
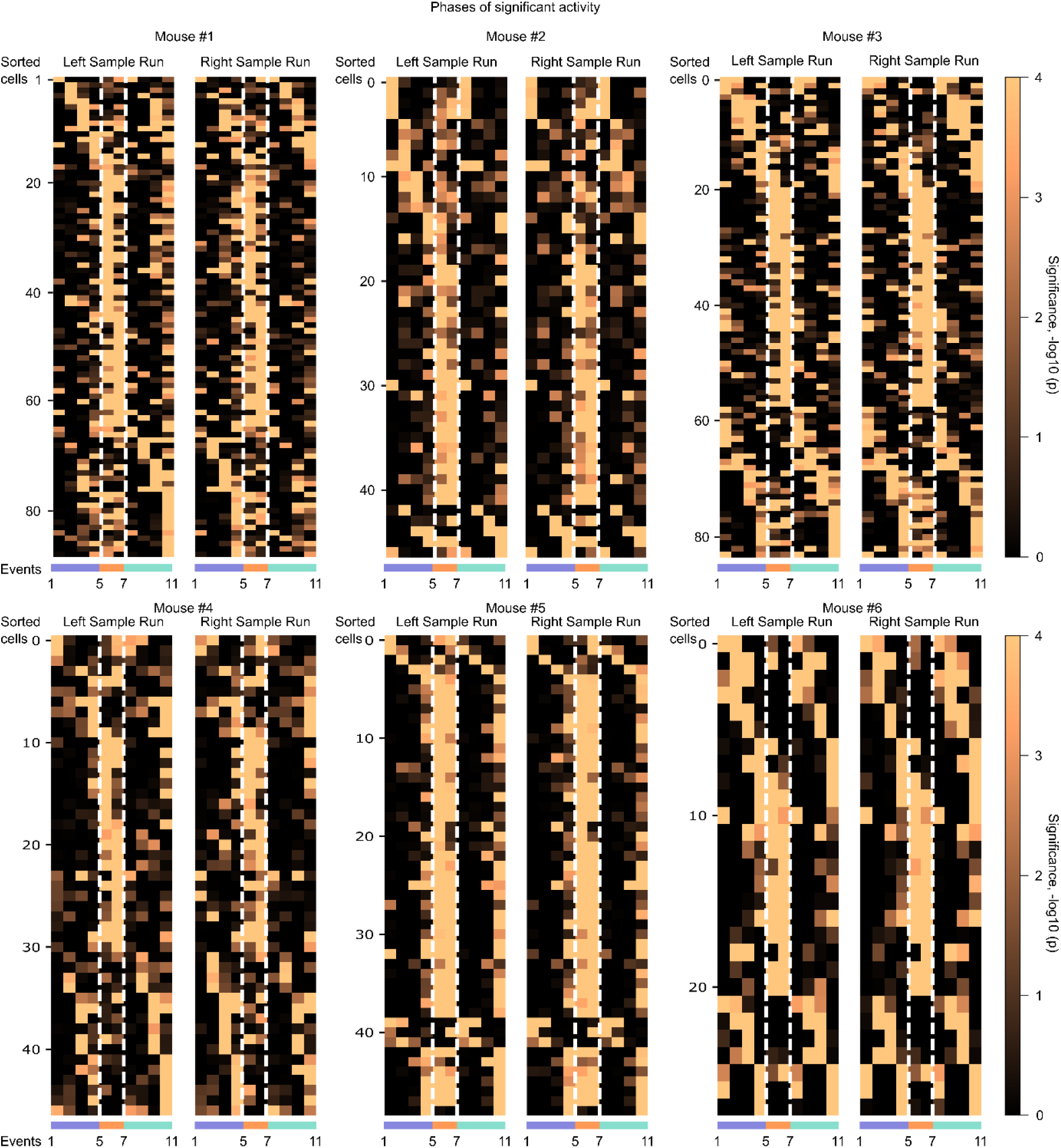
Relative significance of single-cell activity during task phases for all individual mice. Significance level was calculated as -log10(p), where p is the p-value of the Wilcoxon rank sum test, comparing the mean activity from the deconvolved ΔF/F traces for each neuron in a given phase to all other phases. Heatmaps show results separately for left and right sample runs. Single cells were sorted according to the phase with the highest significance level in correct left sample trials. In all mice except for mouse #6, more cells were significantly more active in the maintenance period compared to encoding or retrieval periods (see quantification in Fig. 6f). Note that in some mice, e.g. m#1 and m#3, differences in the neuronal population activity during encoding and retrieval are apparent for left versus right turns. Number of longitudinally recorded neurons: 88 (m1), 46 (m2), 83 (m3), 47 (m4), 49 (m5), 28 (m6).

**Supplementary Figure 9.**
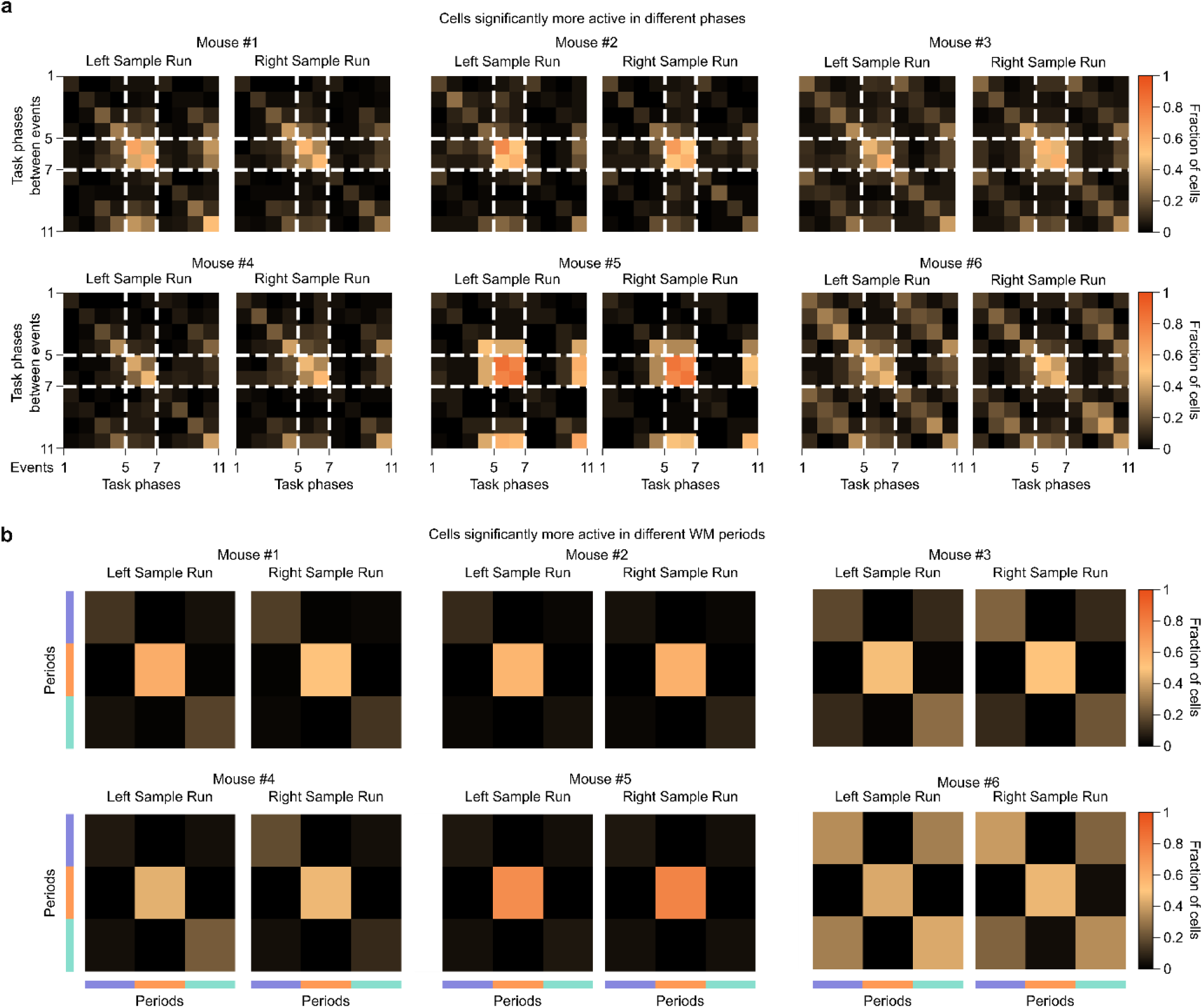
Fraction of cells active in pairs of task phases and task periods for all individual mice. Matrix plots showing the fraction of neurons with significant activity in pairs of task phases (**a**) or task periods (**b**) for each individual mouse. Values on the diagonal represent fraction of neurons significantly more active in the respective phase or period than in other phases/periods. Off-diagonal entries represent the fraction of neurons significantly more active in two different phases/periods simultaneously, compared to other phases/periods. Note the overlap of neuronal fractions that show significant activity in encoding and retrieval periods as well as in corresponding phases during these periods. Coloured bars in (b) correspond to task periods (purple -encoding, orange maintenance, cyan - retrieval). Except for mouse #6, the fraction of neurons active in the maintenance period is higher than in encoding or retrieval periods.

**Supplementary Figure 10.**
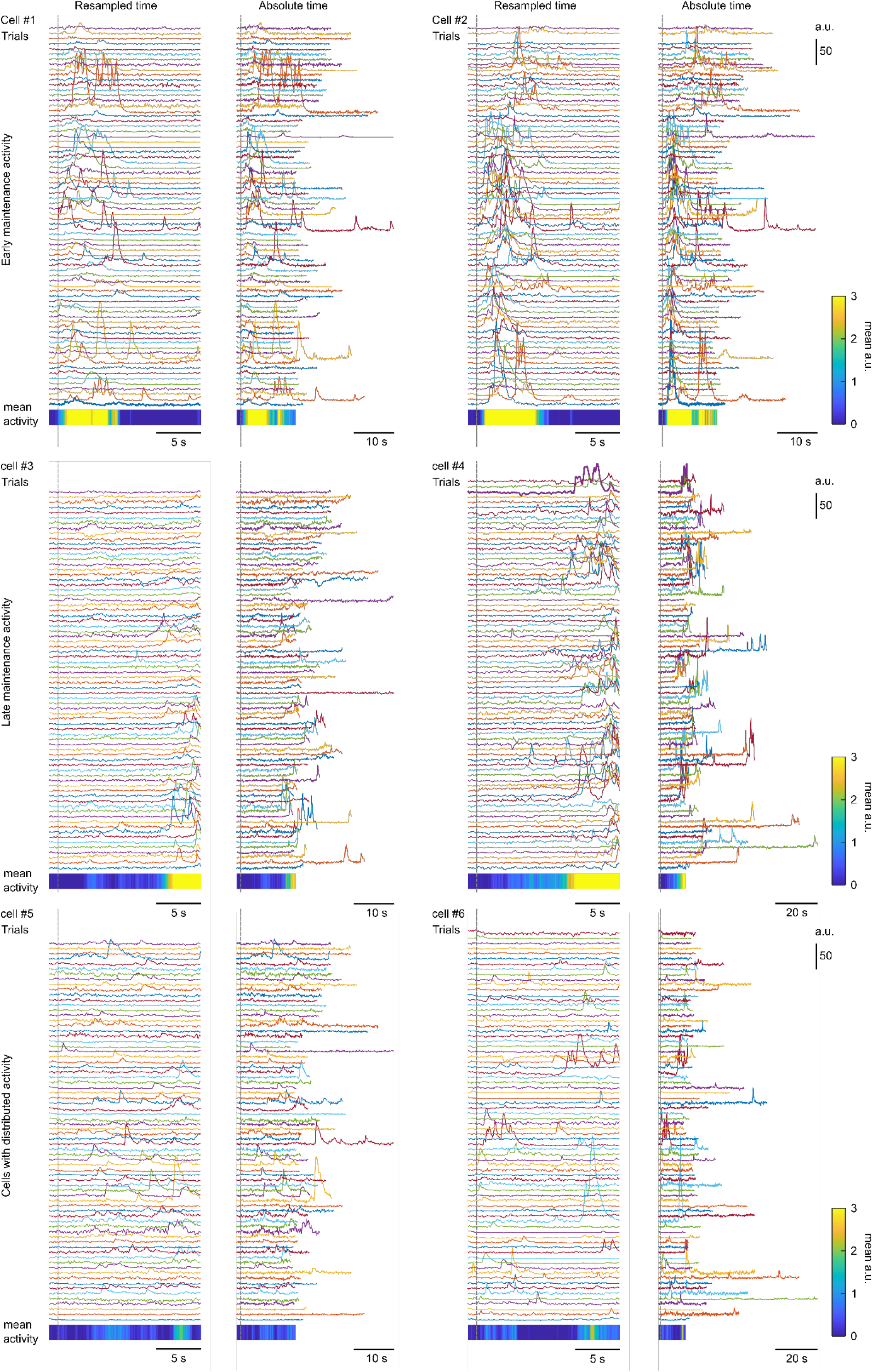
Examples of cells displaying elevated activity during the maintenance period. For each cell, the raw output signal is plotted in arbitrary units (Similar to Fig. 6c left; Methods). The first column shows the cellular activity traces during the whole maintenance period (between events 5 and 7) resampled to the normalized time (median of the 5-7 duration over all trials of all sessions and mice). The heat map shows the mean activity. The second column shows the activity traces during maintenance in absolute time. Since the mice are freely moving, the time spent in the start box varies in some trial although all doors of the maze are open allowing mice to proceed to the choice run. Top row shows the example cells displaying activity in early maintenance period, middle row shows cells active in the late maintenance period. The bottom row shows the cells, which were identified as delay cells, but the time point of their highest activity is not consistent from trial to trial.

**Supplementary Figure 11.**
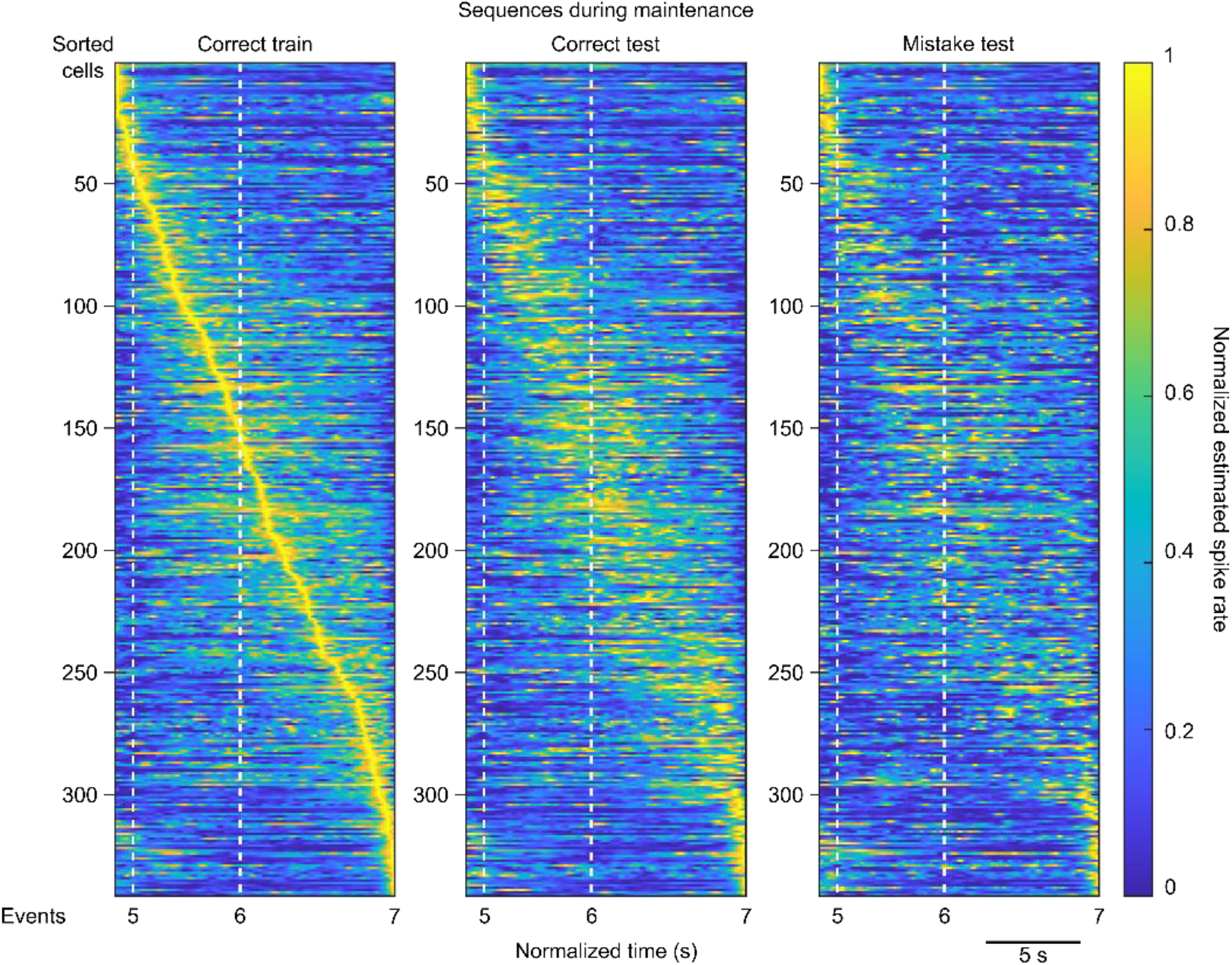
Sequential activity of mPFC→dmSt projection neurons during the maintenance period. Normalized deconvolved neuronal signals during the two phases of the maintenance period pooled for the miniscope data from all 6 mice (n = 341 neurons in total). Left: Mean signals for a randomly selected half of correct trials used as training set. Middle: Mean signals for the remaining test set of correct trials. Right: Mean signals for mistake trials. In all plots, neurons are sorted according to the signal peak times in the training data.

**Supplementary Figure 12.**
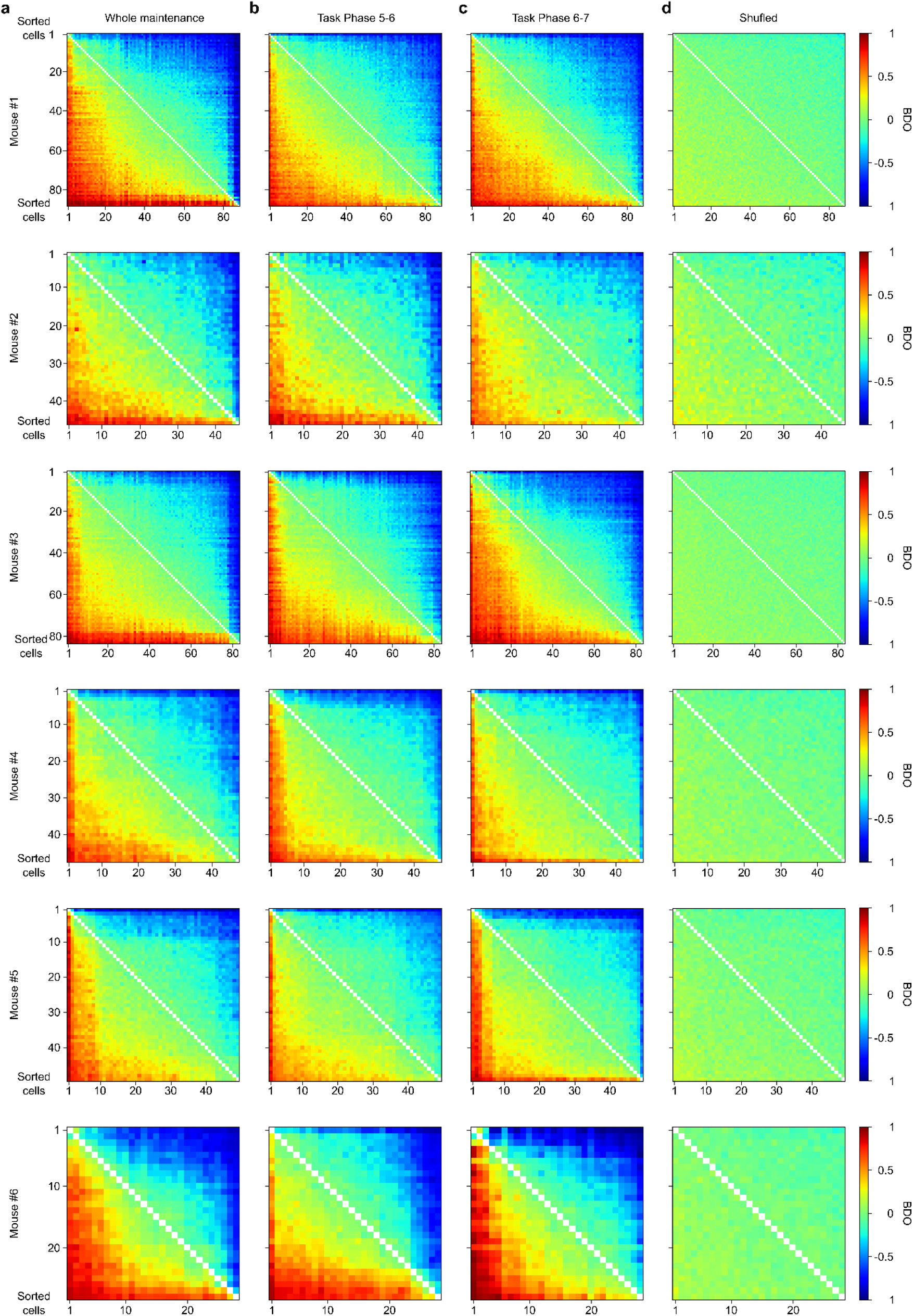
Binary Directed Orderability matrixes for all individual mice. Each row shows the analysis for one mouse. (**a**) For the whole maintenance period. (**b**) For the first maintenance phase between events 5 and 6. (**c**) For the second maintenance phase between events 6 and 7. (**d**) For shuffled data as control. For each plot, the neurons (and thus rows and columns) are sorted according to the neuron’s average BDO of the BDO corresponding to that plot, which, in general, results in a different order of neurons for different plots.

**Supplementary Figure 13.**
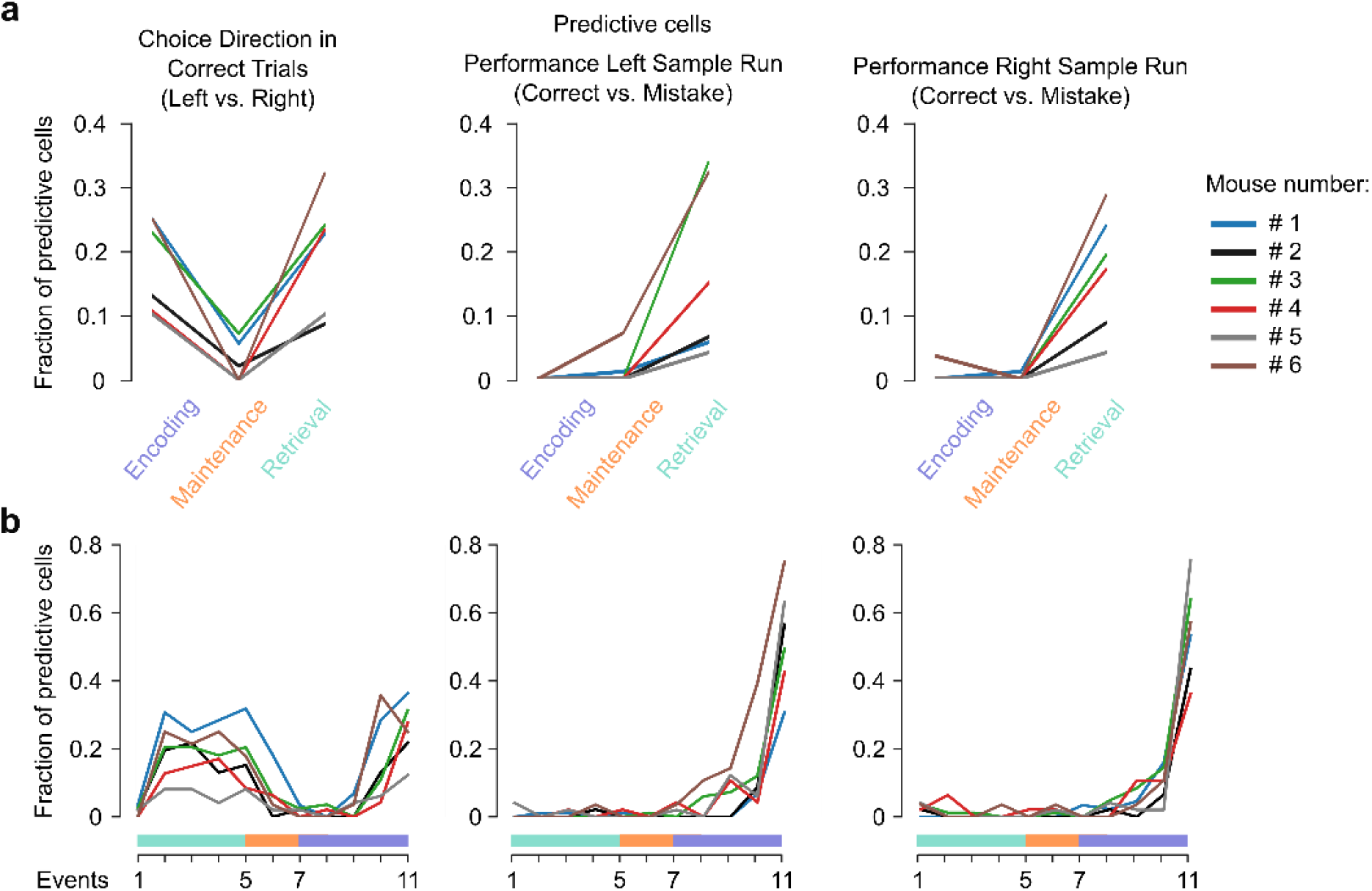
Predictive power of mPFC→dmStr neuronal populations to classify left vs. right turning direction and correct vs. mistake choices across trial task time. (**a**) Fraction of cells predictive for choice direction (left vs. right turn and for performance (correct vs. mistake trials; separated for trials with left and right sample arm) in the different task periods. For definition of predictive neurons see Methods. (**b**) Same as for (a) but for all 10 trial phases. Each coloured line represents an individual mouse. For the turning direction, the fraction of cells predictive for encoding or retrieval periods is higher than for the maintenance period, which might correspond to the unique sub-population of cells active either in right or left maze-arm.

**Supplementary Figure 14.**
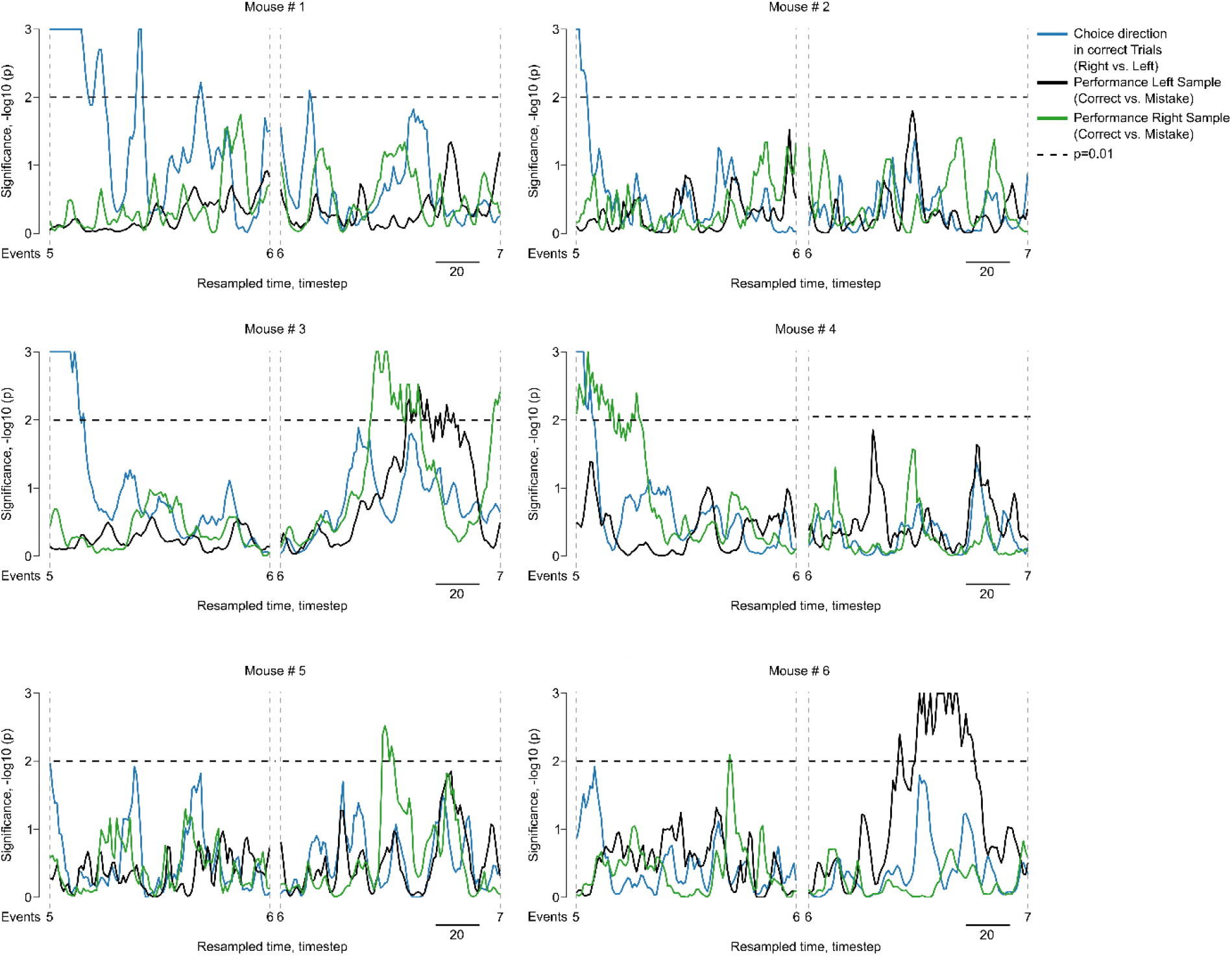
Predictive power of mPFC→dmStr neuronal populations during the two phases of the maintenance period. We tested for population encoding of choice direction (left vs. right turn; blue traces) and of performance (correct vs. mistake trials; separated for trials with left sample arm [black traces] and right sample arm [green traces]). The analysis is based on the L2 norm distance between population vectors for the two considered conditions at each time bin of resampled trial time. We tested against data with shuffled condition labels (Methods). Dashed horizontal lines represent -log10(p)=2, thus indicating chance level of p = 0.01. Each subpanel shows the analysis for an individual mouse. Note that mouse #3 and #6 transiently reach significance during the second phase of the maintenance period for encoding performance when an upcoming contralateral (right) turn is correct. All traces during each maintenance phase have been resampled to 100 time bins for this analysis.

## REFERENCES

1. Curtis, C. E. & D’esposito, M. The effects of prefrontal lesions on working memory performance and theory. Cogn. Affect. Behav. Neurosci. 4, 528–539 (2004).

2. Rossi, M. A. et al. Prefrontal cortical mechanisms underlying delayed alternation in mice. J. Neurophysiol. 108, 1211–1222 (2012).

3. Lewis, D. A. & Gonzalez-Burgos, G. Intrinsic excitatory connections in the prefrontal cortex and the pathophysiology of schizophrenia. Brain Res. Bull. 52, 309–317 (2000).

4. Minzenberg, M. J., Laird, A. R., Thelen, S., Carter, C. S. & Glahn, D. C. Meta-analysis of 41 Functional Neuroimaging Studies of Executive Function in Schizophrenia. Arch. Gen. Psychiatry 66, 811–822 (2009).

5. Perlstein, W. M., Carter, C. S., Noll, D. C. & Cohen, J. D. Relation of Prefrontal Cortex Dysfunction to Working Memory and Symptoms in Schizophrenia. Am. J. Psychiatry 158, 1105–1113 (2001).

6. Cassaday, H. J., Nelson, A. J. D. & Pezze, M. A. From attention to memory along the dorsal-ventral axis of the medial prefrontal cortex: some methodological considerations. Front. Syst. Neurosci. 8, 160 (2014).

7. Spellman, T. et al. Hippocampal-prefrontal input supports spatial encoding in working memory. Nature 522, 309–314 (2015).

8. Bolkan, S. S. et al. Thalamic projections sustain prefrontal activity during working memory maintenance. Nat. Neurosci. 20, 987–996 (2017).

9. Li, Z. et al. The Corticostriatal Adenosine A2A Receptor Controls Maintenance and Retrieval of Spatial Working Memory. Biol. Psychiatry 83, 530–541 (2018).

10. Pan, W. X., Mao, T. & Dudman, J. T. Inputs to the Dorsal Striatum of the Mouse Reflect the Parallel Circuit Architecture of the Forebrain. Front. Neuroanat. 4, (2010).

11. Tervo, D. G. R. et al. A Designer AAV Variant Permits Efficient Retrograde Access to Projection Neurons. Neuron 92, 372–382 (2016).

12. Kang, S. et al. Down-regulation of dorsal striatal RhoA activity and impairment of working memory in middle-aged rats. Neurobiol. Learn. Mem. 103, 3–10 (2013).

13. Moussa, R., Poucet, B., Amalric, M. & Sargolini, F. Contributions of dorsal striatal subregions to spatial alternation behavior. Learn. Mem. 18, 444–451 (2011).

14. Akhlaghpour, H. et al. Dissociated sequential activity and stimulus encoding in the dorsomedial striatum during spatial working memory. eLife 5, e19507 (2016).

15. Park, J. C., Bae, J. W., Kim, J. & Jung, M. W. Dynamically changing neuronal activity supporting working memory for predictable and unpredictable durations. Sci. Rep. 9, 1–10 (2019).

16. Yang, S.-T., Shi, Y., Wang, Q., Peng, J.-Y. & Li, B.-M. Neuronal representation of working memory in the medial prefrontal cortex of rats. Mol. Brain 7, 61 (2014).

17. Bygrave, A. M. et al. Knockout of NMDA-receptors from parvalbumin interneurons sensitizes to schizophrenia-related deficits induced by MK-801. Transl. Psychiatry 6, e778 (2016).

18. Kellendonk, C., Simpson, E. H. & Kandel, E. R. Modeling cognitive endophenotypes of schizophrenia in mice. Trends Neurosci. 32, 347–358 (2009).

19. Sigurdsson, T., Stark, K. L., Karayiorgou, M., Gogos, J. A. & Gordon, J. A. Impaired hippocampal–prefrontal synchrony in a genetic mouse model of schizophrenia. Nature 464, 763–767 (2010).

20. Galizio, M. et al. Effects of NMDA antagonist dizocilpine (MK-801) are modulated by the number of distractor stimuli in the rodent odor span task of working memory. Neurobiol. Learn. Mem. 161, 51–56 (2019).

21. Hurtubise, J. L. et al. MK-801-induced impairments on the trial-unique, delayed nonmatching-to-location task in rats: effects of acute sodium nitroprusside. Psychopharmacology (Berl.) 234, 211–222 (2017).

22. van der Staay, F. J., Rutten, K., Erb, C. & Blokland, A. Effects of the cognition impairer MK-801 on learning and memory in mice and rats. Behav. Brain Res. 220, 215–229 (2011).

23. Yang, Y. & Mailman, R. B. Strategic neuronal encoding in medial prefrontal cortex of spatial working memory in the T-maze. Behav. Brain Res. 343, 50–60 (2018).

24. Cui, G. et al. Concurrent activation of striatal direct and indirect pathways during action initiation. Nature 494, 238–242 (2013).

25. Han, X. et al. A high-light sensitivity optical neural silencer: development and application to optogenetic control of non-human primate cortex. Front. Syst. Neurosci. 5, 18 (2011).

26. Babl, S. S., Rummell, B. P. & Sigurdsson, T. The Spatial Extent of Optogenetic Silencing in Transgenic Mice Expressing Channelrhodopsin in Inhibitory Interneurons. Cell Rep. 29, 1381–1395.e4 (2019).

27. Bajo, V. M. et al. Silencing cortical activity during sound-localization training impairs auditory perceptual learning. Nat. Commun. 10, 3075 (2019).

28. Thuault, S. J. et al. Prefrontal Cortex HCN1 Channels Enable Intrinsic Persistent Neural Firing and Executive Memory Function. J. Neurosci. 33, 13583–13599 (2013).

29. Wang, M. et al. α2A-Adrenoceptors Strengthen Working Memory Networks by Inhibiting cAMP-HCN Channel Signaling in Prefrontal Cortex. Cell 129, 397–410 (2007).

30. Wang, M. et al. Neuronal basis of age-related working memory decline. Nature 476, 210–213 (2011).

31. McClure, K. J. et al. Discovery of a novel series of selective HCN1 blockers. Bioorg. Med. Chem. Lett. 21, 5197–5201 (2011).

32. Flusberg, B. A. et al. High-speed, miniaturized fluorescence microscopy in freely moving mice. Nat. Methods 5, 935–938 (2008).

33. Grewe, B. F. et al. Neural ensemble dynamics underlying a long-term associative memory. Nature 543, 670–675 (2017).

34. Rupprecht, P. et al. A deep learning toolbox for noise-optimized, generalized spike inference from calcium imaging data. bioRxiv 2020.08.31.272450 (2020) doi:10.1101/2020.08.31.272450.

35. Jones, M. W. & Wilson, M. A. Theta Rhythms Coordinate Hippocampal–Prefrontal Interactions in a Spatial Memory Task. PLOS Biol. 3, e402 (2005).

36. Liu, T., Bai, W., Xia, M. & Tian, X. Directional hippocampal-prefrontal interactions during working memory. Behav. Brain Res. 338, 1–8 (2018).

37. Dolleman-van der Weel, M. J. et al. The nucleus reuniens of the thalamus sits at the nexus of a hippocampus and medial prefrontal cortex circuit enabling memory and behavior. Learn. Mem. 26, 191–205 (2019).

38. Jayachandran, M. et al. Prefrontal Pathways Provide Top-Down Control of Memory for Sequences of Events. Cell Rep. 28, 640–654.e6 (2019).

39. Parnaudeau, S., Bolkan, S. S. & Kellendonk, C. The Mediodorsal Thalamus: An Essential Partner of the Prefrontal Cortex for Cognition. Biol. Psychiatry 83, 648–656 (2018).

40. Miller, R. L. A., Francoeur, M. J., Gibson, B. M. & Mair, R. G. Mediodorsal Thalamic Neurons Mirror the Activity of Medial Prefrontal Neurons Responding to Movement and Reinforcement during a Dynamic DNMTP Task. eneuro 4, ENEURO.0196-17.2017 (2017).

41. Yartsev, M. M., Hanks, T. D., Yoon, A. M. & Brody, C. D. Causal contribution and dynamical encoding in the striatum during evidence accumulation. eLife 7, e34929 (2018).

42. Bari, B. A. et al. Stable Representations of Decision Variables for Flexible Behavior. Neuron 103, 922–933.e7 (2019).

43. Emmons, E. B. et al. Rodent Medial Frontal Control of Temporal Processing in the Dorsomedial Striatum. J. Neurosci. 37, 8718–8733 (2017).

44. Mello, G. B. M., Soares, S. & Paton, J. J. A Scalable Population Code for Time in the Striatum. Curr. Biol. 25, 1113–1122 (2015).

45. Duvarci, S. et al. Impaired recruitment of dopamine neurons during working memory in mice with striatal D2 receptor overexpression. Nat. Commun. 9, 1–13 (2018).

46. Hahn, B., Robinson, B. M., Leonard, C. J., Luck, S. J. & Gold, J. M. Posterior Parietal Cortex Dysfunction Is Central to Working Memory Storage and Broad Cognitive Deficits in Schizophrenia. J. Neurosci. 38, 8378–8387 (2018).

47. Scott, G. A., Roebuck, A. J., Greba, Q. & Howland, J. G. Performance of the trial-unique, delayed non-matching-to-location (TUNL) task depends on AMPA/Kainate, but not NMDA, ionotropic glutamate receptors in the rat posterior parietal cortex. Neurobiol. Learn. Mem. 159, 16–23 (2019).

48. Sych, Y., Chernysheva, M., Sumanovski, L. T. & Helmchen, F. High-density multi-fiber photometry for studying large-scale brain circuit dynamics. Nat. Methods 16, 553–560 (2019).

49. Stubbendorff, C., Molano-Mazon, M., Young, A. M. J. & Gerdjikov, T. V. Synchronization in the prefrontal–striatal circuit tracks behavioural choice in a go–no-go task in rats. Eur. J. Neurosci. 49, 701–711 (2019).

50. Terra, H. et al. Prefrontal Cortical Projection Neurons Targeting Dorsomedial Striatum Control Behavioral Inhibition. Curr. Biol. 30, 4188–4200.e5 (2020).

51. Homayoun, H. & Moghaddam, B. NMDA Receptor Hypofunction Produces Opposite Effects on Prefrontal Cortex Interneurons and Pyramidal Neurons. J. Neurosci. 27, 11496–11500 (2007).

52. Jackson, M. E., Homayoun, H. & Moghaddam, B. NMDA receptor hypofunction produces concomitant firing rate potentiation and burst activity reduction in the prefrontal cortex. Proc. Natl. Acad. Sci. 101, 8467–8472 (2004).

53. Paspalas, C. D., Wang, M. & Arnsten, A. F. T. Constellation of HCN Channels and cAMP Regulating Proteins in Dendritic Spines of the Primate Prefrontal Cortex: Potential Substrate for Working Memory Deficits in Schizophrenia. Cereb. Cortex N. Y. NY 23, 1643–1654 (2013).

54. Tokay, T. et al. HCN1 channels constrain DHPG-induced LTD at hippocampal Schaffer collateral-CA1 synapses. Learn. Mem. 16, 769–776 (2009).

